# Traveling wave chemotaxis of neutrophil-like HL-60 cells

**DOI:** 10.1101/2024.06.16.598630

**Authors:** Motohiko Ishida, Masahito Uwamichi, Akihiko Nakajima, Satoshi Sawai

## Abstract

The question of how changes in chemoattractant concentration translate into the chemotactic response of immune cells serves as a paradigm for the quantitative understanding of how cells perceive and process temporal and spatial information. Here, using a microfluidic approach, we analyzed the migration of neutrophil-like HL-60 cells to a traveling wave of the chemoattractants fMLP and leukotriene B4 (LTB4). We found that under a pulsatile wave that travels at a speed of 95 and 170 µm/min, cells move forward in the front of the wave but slow down and randomly orient at the back due to temporal decrease in the attractant concentration. Under a slower wave, cells re-orient and migrate at the back of the wave; thus, cell displacement is canceled out or even becomes negative as cells chase the receding wave. FRET-based analysis indicated that these patterns of movement correlated well with spatiotemporal changes in Cdc42 activity. Furthermore, pharmacological perturbations suggested that migration in front of the wave depends on Cdc42, whereas that in the back of the wave depends more on PI3K/Rac and ROCK. These results suggest that pulsatile attractant waves may recruit or disperse neutrophils, depending on their speed and degree of cell polarization.

**Significance statement:**

- The way neutrophil chemotaxis is directed by attractants is thought to depend on temporal and spatial concentration changes, however the response to a transient pulsatile stimulus has not been well explored.
- A moderately fast traveling wave of fMLP and leukotrieneB4 (LTB4) directs unidirectional cell migration. Under slow waves, cells respond to the back of the wave; thus, cell displacement is canceled out. Cdc42, PI3K, Rac, and ROCK contributed differently to responses at the front and back of the wave.
- These findings suggest that traveling waves of attractant can guide immune cell recruitment and dispersal.

## INTRODUCTION

The movements of fast-migrating cells, such as the amoeba *Dictyostelium* and neutrophils, are driven by F-actin-based crawling motility that is guided and oriented by selective cues. In general, chemotaxis is thought to be mediated by a ‘gradient-sensing’ response in which differences in receptor occupancy along an attractant gradient are translated intracellularly to form an expanding leading edge at one end of the cell and a contracting trailing edge at the other(Tranquillo and Lauffenburger, 1986; Devreotes and Zigmond, 1988). However, the mechanism by which cells perceive and respond to transient and dynamic cues remains unclear. Attractant degradation can give rise to a self-generating gradient, whose spatial profile changes as the cells move(Tweedy *et al*., 2016). As the cells migrate and invade regions with high levels of attractant, a gradient is formed that further induces cell movement. This feedback reinforces each other and generates a moving wavefront. In addition to such degradation, if the attractant is self-amplified and relayed, that is, there is attractant-induced synthesis or release of the attractant, it may give rise to a pulsatile wave(Goldbeter, 1996). An example of such a stimulus is extracellular cAMP, which relays between aggregating *Dictyostelium* cells in the form of propagating waves(Tomchik and Devreotes, 1981; Dormann *et al*., 2000; Sawai *et al*., 2005). While it is still unclear whether similar pulsatile waves exist outside *Dictyostelids*, neutrophils are also known to swarm to a site of inflammation by moving towards leukotriene B4 (LTB4), which is secreted by the neutrophils themselves(Lämmermann *et al*., 2013; Tamás *et al*., 2023). While such a signal relay provides an advantage in terms of the distance at which a signal exerts its influence, cells can be met with the complex task of reading out a transient stimulus composed of a rising phase (wavefront) and an attenuating phase (waveback) that can induce movements with contradictory orientations. From an information processing point of view, this poses the question of what spatial and temporal components of a propagating stimulus are read out by cells to determine their orientation.

In *Dictyostelium* cell aggregation, the long-held notion is that cells orient themselves by sensing a gradient in the wavefront. As cells do not re-orient during waveback, it is expected that there will be a period when cells are refractory or desensitized(Geiger *et al*., 2003). However, when *Dictyostelium* cells rapidly re-orient attractant gradients from a glass needle, a plekstrin homology (PH)-domain-containing protein that serves as an indicator of leading edge formation quickly relocates to the membrane region closest to the needle (Jin *et al*., 2000), indicating an apparent lack of refractoriness. An alternative explanation is the first-hit mechanism(Haston and Wilkinson, 1988; Rappel *et al*., 2002). According to this scheme, the leading-edge response occurs only in the region that is first exposed to the attractant. This requires a fast-spreading inhibitory signal that laterally suppresses the other end before the stimulus arrives and therefore assumes some type of winner-takes-all mechanism(Byrne *et al*., 2014). The origin of such long-range inhibition, if any, is not yet clear, and may be mediated by molecular diffusion (Rappel and Edelstein-Keshet, 2017) or the spread of membrane tension(De Belly *et al*., 2023). Furthermore, the transient and stationary responses are influenced by the memory of cell polarity(Skoge *et al*., 2014; Prentice-Mott *et al*., 2016). Once primed with wavefront stimulation, the established cell polarity is maintained even in the absence of an attractant gradient, as long as its mean concentration remains elevated(Skoge *et al*., 2014). Similarly, an increase in cell velocity has been observed under repetitive uniform stimuli (Varnum et al., 1985) as well as under periodic traveling wave stimuli(Nakajima *et al*., 2016). When polarity persists, cells may re-orient simply by guiding pre-existing leading edge - a mode of chemotaxis known as ‘steering’(Yang *et al*., 2016; Boujemaa-Paterski *et al*., 2017). This could be accomplished by undulating the leading edge towards the lateral sides, which could be biased in one way or the other by the difference in the attraction concentration. This may again be mediated by spatial gradient perception or active sensing of temporal changes, but at a much smaller scale of membrane protrusions(Gerisch *et al*., 1975; Devreotes and Zigmond, 1988; Town and Weiner, 2023).

The quest to understand temporal and spatial perception has also developed around neutrophil chemotaxis, with interesting parallels. A pioneering work by Michael Vicker et al. (Vicker *et al*., 1986) utilized a low-temperature condition to keep the cells immotile, while establishing a gradient of the chemoattractant N-formyl-methionyl-leucyl-phenylalanine (fMLP). This work suggested that for neutrophils to exhibit directional bias, the concentration of chemoattractants needs to increase over time, and thus they may not respond to a spatial gradient. However, this suggestion has been met with the criticism that the data can also be interpreted using gradient perception(Lauffenburger *et al*., 1987). A later study by Albrecht and Petty (Albrecht and Petty, 1998) used a flow chamber to introduce a spatially homogeneous stepwise shift in fMLP concentrations. This work indicates that neutrophils reverse their cell orientation when provided with a temporally decreasing concentration of fMLP. Using two flow pumps to form a moving gradient, Ebrahimzadeh et al. showed that human neutrophil movements are inhibited when the gradients are moved in a direction opposite to that of cell migration to decrease the fMLP concentration over time (Ebrahimzadeh, et al Braide, 2000). Furthermore, a recent study by Irimia et al. devised a system in which the position of the gradient could be updated in real-time with reference to the cell position(Aranyosi *et al*., 2015). They found that positive temporal gradients are necessary for the directional movement of human neutrophils(Aranyosi *et al*., 2015). This observation is also supported by the fact that neutrophil migration no longer persists under a stabilized gradient of fMLP, (Chandrasekaran *et al*., 2017) as well as the chemokines CCL19 and SDF1(Petrie Aronin *et al*., 2017). As for periodic stimuli, Soll et al. realized continuously varying periodic stimuli by employing a flow chamber and two programmable pumps and found remarkable similarity in the response displayed by neutrophils and *Dictyostelium*(Geiger *et al*., 2003). These oscillatory stimuli had a 7 min period and the amplitude was two orders of magnitude: 10–1000 nM cAMP for *Dictyostelium* and 1–100 nM fMLP for neutrophils. In the rising phase, the cells formed a single leading edge, and its velocity increased approximately 2- fold which peaked slightly before the top of the stimuli and fell quickly to the pre-stimulus level. In the falling phase, contrary to the earlier observation of reorientation(Albrecht and Petty, 1998), the cells were non-polarized, and the velocity remained low at the basal level.

These studies lead to the intriguing hypothesis that in a manner similar to *Dictyostelium* aggregation, neutrophils may be capable of being guided by pulsatile waves of attractants (Geiger *et al*., 2003). This property may be beneficial in vivo when neutrophils are recruited from long distances to a site of inflammation and tissue damage. It may also be that the cessation of neutrophil recruitment during inflammatory resolution is related to the manner in which cells respond to time-varying stimuli(Chandrasekaran *et al*., 2017). To date, whether and, if so, how stimuli in the form of traveling waves orient and guide neutrophil migration has not been directly tested. Analyzing such behaviors should not only provide hints on the migratory repertoires of neutrophils in vivo, but also a deeper understanding of chemotactic sensing and signal perception in time and space(Levchenko and Iglesias, 2002; Wang *et al*., 2014; Petrie Aronin *et al*., 2017; Hadjitheodorou *et al*., 2021). In this study, based on live-cell imaging and a microfluidics approach, we analyzed the chemotaxis of neutrophil-like HL-60 cells in response to a traveling wave stimulus. Our analysis showed that HL-60 cells can orient unidirectionally towards a wave stimulus that travels at intermediate speeds for both fMLP and LTB4. Based on the speeds at which reverse migration at the waveback is suppressed and the pharmacological perturbation that reverses this effect, we discuss possible mechanisms that underlie spatio-temporal stimulus perception and their implications.

## RESULTS

### HL-60 cells exhibit unidirectional migration towards traveling wave stimulus

To address the migration of neutrophil-like HL-60 cells under traveling wave stimuli, we employed a laminar flow-based approach previously used in *Dictyostelium*(Meier *et al*., 2011; Nakajima *et al*., 2014; Nakajima and Sawai, 2016). A bell-shaped gradient of chemoattractant fMLP or LTB4 was generated in a microfluidic chamber with three inlets (Ibidi 80311, µ-Slide III 3in1 uncoated; Figure 1A, left panels). By changing the flow speed from the side channels in a reciprocal manner (Figure 1A, right panel), the bell-shaped profile was continuously displaced in a direction orthogonal to the flow. The flow speed from the center channel was 3 µL/min and that from the side channels varied from 4 to 26 µL/min (Figure 1A, right panel). The shear stress resulting from the flow viscosity (Décavé *et al*., 2003) was approximately 5 × 10^-3^ Pa which is two orders of magnitude less than that required for shear-stress induced motility in HL-60 cells (Makino et al 2005). As the wave swept across the chamber, a cell was first exposed to a temporally increasing gradient in the mean chemoattractant concentration (referred to hereafter as ‘wavefront’), followed by a reverse gradient (‘waveback’) that was temporally decreasing in the mean concentration (Figure 1A, left lower panel). The base and peak stimulus source concentrations were set to 1 and 50 nM for fMLP and 0 and 2 nM for LTB4 so that the wave profile would cross over the dissociation constant; 1-10 nM for the fMLP receptor (Atkinson 1988, Quehenberger 1993, Herzmark 2007) and 0.15 nM for the LTB4 receptor(Yokomizo *et al*., 1997). The length of the wave profile chosen in this study was fixed to *L*p = 668 µm which was defined by a region where the attractant concentration was above 2% of the source concentration (Figure 1B). At the mid-point of the slope, the imposed fMLP gradient was approximately 10% change across the cell diameter (i.e. ∼ 30 pM/µm) which was two orders of magnitude steeper than the threshold steepness known for human neutrophil(Chandrasekaran *et al*., 2017).

**FIGURE 1:**
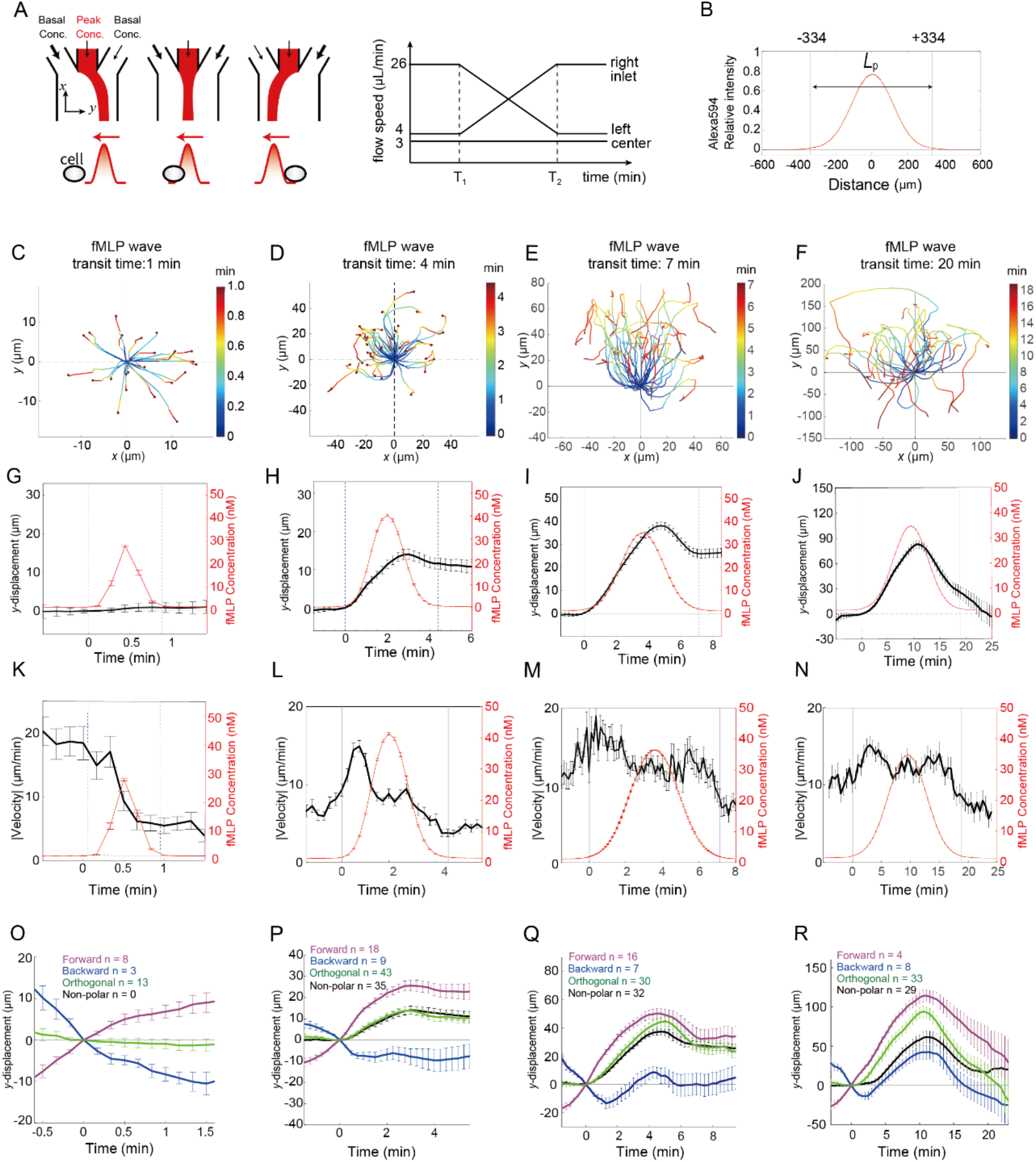
Migratory response of HL-60 cells to a traveling wave of fMLP depends on the propagation speed. (A) Schematics of the wave experiment. Speed of buffer infusion from the two side channels is continuously and reciprocally varied so that the laminar flow from the center channel (red) is smoothly displaced in time. The buffer in the side channels contains 1 nM fMLP, and that in the center contains 50 nM fMLP and 0.4 µg/mL Alexa 594. The y-axis is orthogonal to the flow direction. The ramp time T2-T1 determines the propagation speed. (B) The spatial profile of the wave stimulus. (C-F) Cell centroid trajectories for wave transit time of 1 min (C), 4 min (D), 7 min (E) and 20 min (F). The origin is set to the initial position at the time of stimulus arrival (t = 0). (G-J) The average centroid displacement in y-direction (black) in reference to the fMLP level (red) for wave transit time of 1 min (G), 4 min (H), 7 min (I) and 20 min (J). The average absolute speed of centroids for wave transit time of 1 min (K), 4 min (L), 7 min (M), and 20 min (N). n = 31 (G and K), 105 (H and L), 85 (I and M), and 87 cells (J and N). (O-R) The mean y-displacement of cell centroids based on the initial cell orientation for wave transit time of 1 min (O), 4 min (P), 7 min (Q) and 20 min (R). Forward (magenta), Backward (blue), Orthogonal (green), and Non-polar (black); see Supplemental Figure S2A for notiation).

Figure 1, C-F, show cell trajectories during the passage of a single fMLP wave. Under a 1 min ‘transit time wave (wave speed: *L*p /1 min = 6.7 × 10^2^ µm/min), HL-60 cells showed on average no (re-)orientation during the stimulus (Figure 1, C and G). Under a slower wave of 4 min transit time (wave speed: *L*p / 4 min = 1.7 × 10^2^ µm/min), cells moved in the direction opposite to that of the wave propagation in the wavefront followed by almost no displacement in the waveback (Figure 1, D and H). In both conditions, the maximum average velocity was about 15–20 µm/min, which dropped to 10 µm/min after 0.5–1 min from the stimulus arrival (Figure 1, K and L). The average and median of the absolute value of the chemotactic index for a 4 min wave remains relatively small in the first half of the wavefront and even more so in the waveback (Supplemental Figure S1A; -Ttransit/4 and Ttransit/4). Thus, the unidirectionality under the 4 min wave can be attributed to decreased cell velocity and cell orientation being slightly more random at the waveback (Supplemental Figure S1A; Ttransit/4) (Figure 1L). For a transit time of 7 min (wave speed: *L*p / 7 min = 9.5 × 10 µm/min), cells climbed up the wavefront then reversed their direction approximately 2 min after the passage of the stimulus peak (Figure 1, E and I). The behavior was similar for a slower wave of 20 min transit time (wave speed: *L*p / 20 min = 3.3 × 10 µm/min), except that reverse movement persisted and the net displacement was even slightly negative (Figure 1, F and J) owing to cells chasing the waveback thus self-extending the stimulus duration. For both 7- and 20 min waves, the average velocity remained elevated above 10 µm/min, which slightly decreased near the peak (Figure 1, M and N). Chemotactic index indicated clear cell (re-)orientation taking the value close to +1 in the wavefront then flipping to -1 in the waveback (Supplemental Figure S1, B and C).

At the individual cell level, our data showed that the ability of HL-60 cells to orient in the wavefront depended strongly on the cell orientation prior to stimulus arrival (Figure 1, O-R; see Supplemental Figure S2A for orientation nomenclature). At all the four transit speeds tested, the cells that migrated in the forward direction maintained their orientation in the wavefront (Figure 1, O-R; magenta). For cells that moved backward before stimulus arrival, almost no change in movement was observed for the 1 min wave, (Figure 1O; blue). Under a 4 min wave, backward-moving cells stalled within 30 s of stimulus arrival and remained there for the duration of our observation (Figure 1P; blue). In the 7- and 20 min waves, backward cells reoriented approximately 1 min after stimulus arrival and migrated forward (Figure 1, Q and R; blue). Cells that were first moving in the orthogonal direction were also oriented towards the wavefront for the 4-, 7-, and 20 min wave but not for the 1 min wave (Figure 1, O-R; green). These results show that, in the wavefront too, cell re-orientation occurred only at slow wave speeds. As for cells that were ‘non-polar’ and thus not migrating in a particular direction before the stimulus, they were all directed by 4, 7, and 20 min waves (Figure 1, P-R; black). Owing to small sampling, our 1 min wave data did not contain cells that were ‘non-polar’ prior to the stimulus (Figure 1O), which is the reason the average velocity was already elevated before the wave arrived (Figure 1K). In contrast to the wavefront, cell movement in the waveback did not show a lasting influence of pre-stimulus orientation. At fast waves (1- and 4 min transit times), there were no clear reverse movement (Figure 1, O and P). In slower waves (7- and 20 min transit time), the cells were re- oriented (Figure 1, Q and R). Regardless of the pre-stimulus cell orientation, there was a slight delay of approximately 1 min between peak arrival and the time the cells began to move backward. In general, neutrophils can re-orient in two ways: by reversing their polarity, or by making a U-turn while maintaining their polarity(Hind *et al*., 2016). We found an almost equal occurrence of these two patterns in the 7 min wave (Supplemental Figure S3). In the 20 min waves, the percentage of cells that made a U-turn increased (Supplemental Figure S3), suggesting that polarity was strengthened under slow waves.

The above results demonstrated that HL-60 cells on average orient and move unidirectionally towards an incoming wave of fMLP that travels at intermediate speeds (95 and 170 µm/min). No stimulus-induced orientation was observed under a fast wave (670 µm/min). In a slow wave (33 µm/min), cells not only moved towards the wavefront but also chased the back of the passing wave and thus the net displacement was close to zero. Next, we tested HL-60 cell migration in a traveling wave of the endogenous chemoattractant LTB4. LTB4 is a secondary chemoattractant released from neutrophils stimulated by end-target chemoattractants such as fMLP(Afonso *et al*., 2012; Lämmermann *et al*., 2013). Secretion and relayed amplification of LTB4 help recruit neutrophils from far distances to the site of infection and tissue damage(Lämmermann *et al*., 2013; Szatmary *et al*., 2017; Hopke *et al*., 2022; Tamás *et al*., 2023). Under the LTB4 wave, the trajectories of HL-60 cells showed less directional bias (Figure 2, A-C) than those under fMLP waves (Figure 1, D-F). Nonetheless, the average displacement showed that for 4 and 7 min LTB4 waves, HL-60 cells migrated in the wavefront but not in the waveback (Figure 2, D and E). The average cell displacement in the waveback under a 7 min wave (Figure 2E) was suppressed more than that under the fMLP wave at the same speed (Figure 1I). For a transit time of 20 min, cells, on average, migrated towards the wavefront and then reverse-migrated at the waveback (Figure 2F). Compared with the fMLP, the velocity response showed a very weak peak (Figure 2, G-I). The average velocity after the passage of the stimulus peak declined from about 20 to 10 µm/min for 4- and 7- min waves (Figure 2, G and H). For the 20 min wave, the average velocity persisted (Figure 2I). The chemotactic index indicated that cell orientation was more broadly distributed with LTB4 (Supplemental Figure S4, A-C) than with fMLP (Supplemental Figure S1, A-C). The pre-stimulus orientation affected the migratory pattern in a manner similar to fMLP, except that the wave speed dependency appeared to be offset for approximately 3 min (Figure 2, J-L); that is, the response of the backward-moving cell to the 7 min LTB4 wave (Figure 2K; blue) did not re- orient in the wavefront and thus appeared more similar to that under the 4 min fMLP wave (Figure 1P). These data indicate that, similar to fMLP, a traveling wave of LTB4 of intermediate speeds (95 and 170 µm/min) supports unidirectional migration of HL-60 cells.

**FIGURE 2:**
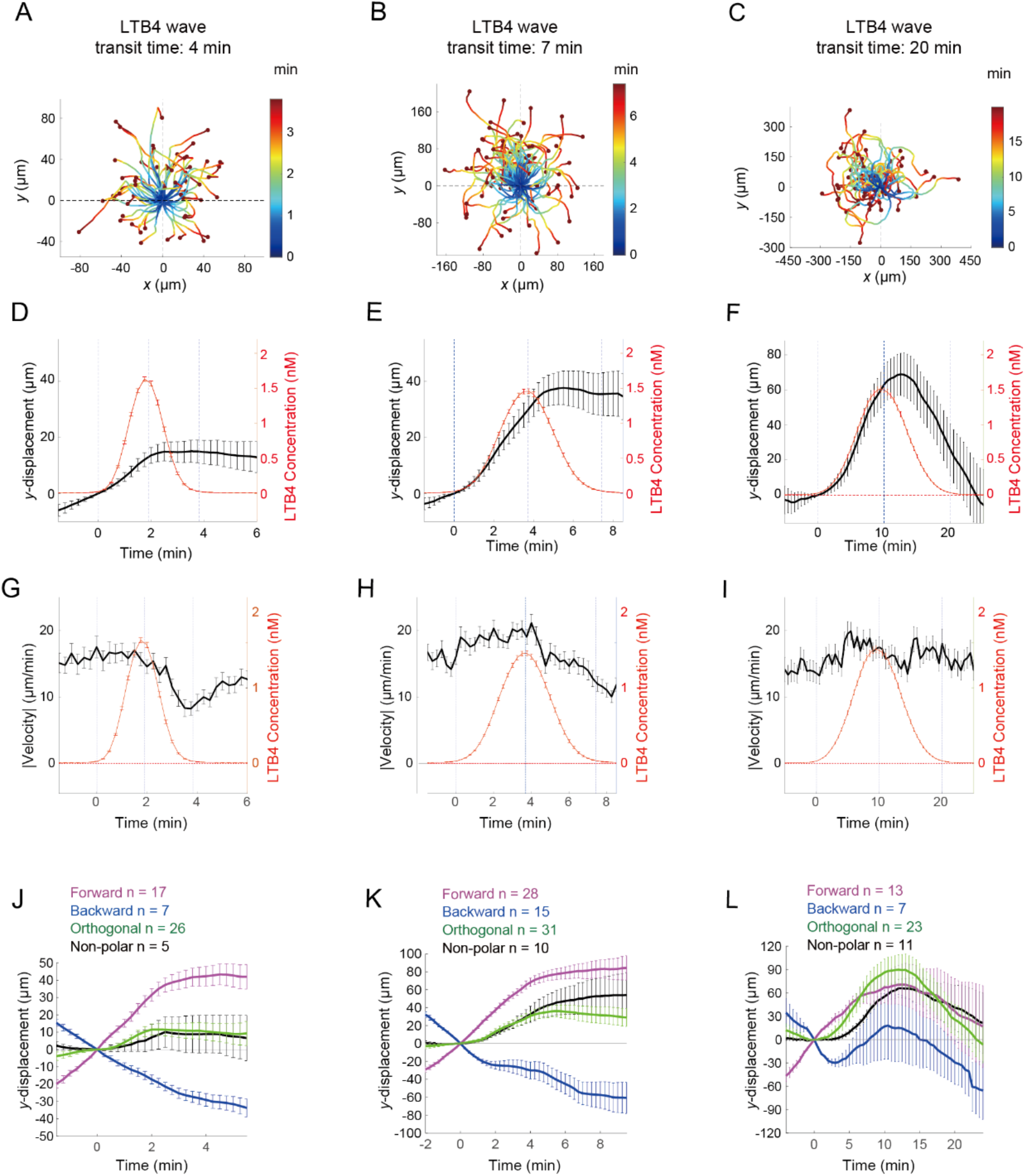
Migratory response of HL-60 cells to LTB4 wave stimulus exhibits similar dependency on the propagation speed. (A-C) Cell centroid trajectories for wave transit time of 4 min (A), 7 min (B) and 20 min (C). The origin is set to the initial position at the time of stimulus arrival (t = 0). (D-F) The average centroid displacement in y-direction (black) in reference to the LTB4 level (red) for wave transit time of 4 min (D), 7 min (E) and 20 min (F). The average absolute speed of centroids for wave transit time of 4 min (G), 7 min (H) and 20 min (I). n = 55 (D and G), 84 (E and H) and 54 cells (F and I). (J-L) The mean y-displacement of cell centroids based on the initial cell orientation for wave transit time of 4 min (J), 7 min (K), and 20 min (L). Forward (magenta), Backward (blue), Orthogonal (green), and Non-polar (black); see Supplemental Figure S2 for notation.

The inability of the HL-60 cells to migrate during the waveback for intermediate-speed waves may be related to the time required for re-orientation. For example, cells may become desensitized after initial exposure to an attractant or may require time to reconstruct cytoskeletal networks. Another possibility is that a temporal decrease in attractant concentration suppresses the chemotactic response regardless of its past gradient history. To explore these and other possibilities, we analyzed responses to an inverted wave stimulus. Here, the order in which cells were exposed to the gradients was flipped, that is, a gradient with a decreasing mean concentration was first followed by a rising reverse gradient (Figure 3A). We set the transit time to 7 min for fMLP and 10 min for LTB4 so that they could be compared with similar asymmetric responses under normal waves. We found that, for fMLP, no directional cell movement was observed on average in the descending gradient (Figure 3B; 0–4 min), in contrast to a large displacement towards the upside in the rising gradient (Figure 3B; 5–10 min). Cell velocity decreased from 10 µm/min to 5 µm/min in the descending gradient which then began to increase in the ascending gradient until it peaked to about 15 µm/min (Figure 3C). Note that at the individual cell level, regardless of whether the cells were moving towards or away from the incoming wave (Figure 3D), the average cell displacement was almost completely flattened within a minute of exposure to the descending gradient (Figure 3D; 0–4 min). This suggests that the decrease in cell velocity was not due to the disparity between the direction of the chemoattractant gradient and the cell polarity. Given that the average cell velocity was still non-zero (Figure 3C) and because the chemotactic index was broadly distributed (Supplemental Figure S4D), it appeared that the descending slope acted to randomize the cell orientation. For LTB4, although there was some displacement towards the descending slope (Figure 3E; 0–5 min), a much larger displacement towards the ascending gradient was observed (Figure 3E; 5–12 min). Notably, the average cell velocity did not drop as much as that under the fMLP, which may be due to the slower speed chosen for the inverted LTB4 wave (Figure 3F). Individual cell tracking indicated that while cells that were initially oriented backward before the gradient arrival continued to move in the same direction, that is, in the ascending direction along the gradient (Figure 3G; blue 0–5 min), those that were initially oriented in other directions ceased to make net movements (Figure 3G; 0–5 min), despite the average absolute velocity being non-zero. Overall, these results suggest that under a temporally descending gradient, the average cell displacement is suppressed mainly due to the randomization of cell orientation, as indicated by the broadly distributed chemotactic index (Supplemental Figure S4E). Under the fMLP, randomization occurred irrespective of the prior cell orientation. Under LTB4, this occurred specifically in cells that did not move in the direction coherent to the imposed attractant gradient, which is consistent with the observation that backward-moving cells continued to move in the same direction during the waveback of the normal LTB4 wave (Figure 2J; t > 2 min and Figure 2K; blue t > 4 min). The fact that this property was only seen under faster waves may be explained by the lack of time required to restructure the cytoskeleton; however, we should note that cells that were initially non-polar were still oriented towards an incoming normal wave (Figures 1Q and 2K black; 0–4 min) but not an inverted wave (Figure 3, D and G; black; 0–4 min), suggesting that the distinction between temporally negative and positive signals is made equally well in polarized and non-polarized cells.

**FIGURE 3:**
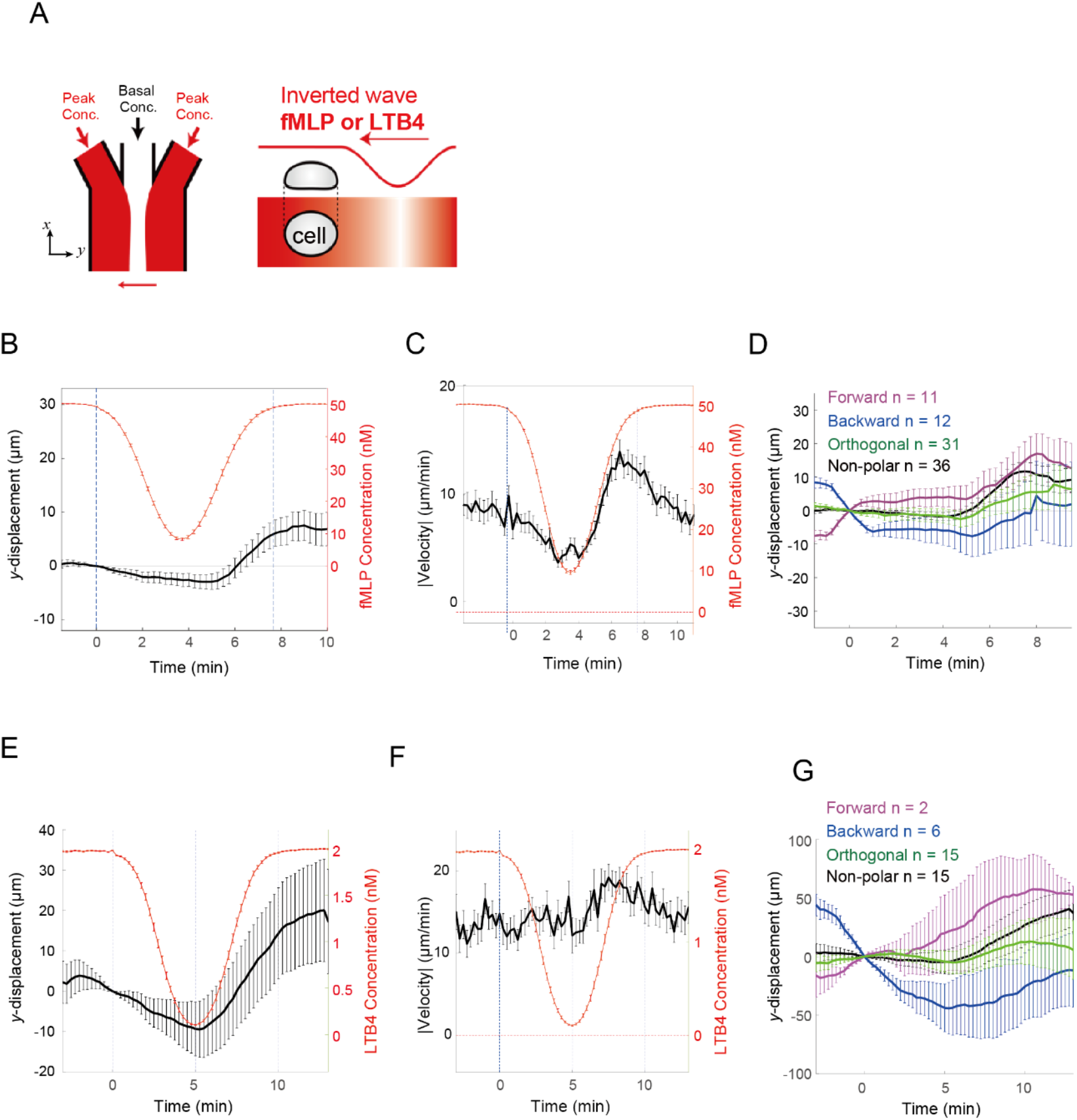
Migratory response of HL-60 cells to inverted wave stimulus. (A) Schematics of the inverted wave experiment. Buffer flow from the two side channels contain either 50 nM fMLP or 2 nM LTB4 in addition to 0.4 µg/mL Alexa594 (red). The flow from the center channel contains 1 nM fMLP or 0 nM LTB4. Flow speed from the side channels is varied according to the scheme in Figure 1A. (B-D) fMLP inverted wave experiments (n = 91 cells). (E-G) LTB4 inverted wave experiments (n = 38 cells). The average y-displacement (B and E), average absolute speed (C and F), and mean y-displacement of cell centroids based on the initial cell orientation (D and G): Forward (magenta), Backward (blue), Orthogonal (green), and Non- polar (black); see Supplemental Figure S2A for notation.

### The distinct migratory response at different wave speeds is correlated with Cdc42 activity

The small GTPase Cdc42 is a key regulator of leading-edge formation in neutrophils(Bell *et al*., 2021; Hadjitheodorou *et al*., 2021). To analyze its relationship with the migratory response under the wave stimulation, we employed fluorescence resonance energy transfer (FRET) analysis using a HL-60 cell line stably expressing the biosensor RaichuEV-Cdc42-CAAX(Komatsu *et al*., 2011).

The spatial difference in the FRET signals of individual cells was obtained by splitting the cell mask into two and quantifying the average. The cell mask was partitioned in two ways: one based on the wave direction, that is, the y-axis in the reference frame ([y+] side, [y-] side; Supplemental Figure S1B) and the other based on the cell polarity axis, as defined by the cell velocity vector (cell front vs. cell rear; Supplemental Figure S1C). Under a 1 min fMLP wave, Cdc42 activity rose and peaked at about 0.5 min then fell below the pre-stimulus level before gradually recovering the basal activity in the following 1 to 2 min time window (Figure 4A). We compared the weighted average <FRET> y+/y- between the side facing the incoming wave ([y+] side) and the opposite side ([y-] side; Supplemental Figure S2B), and the transient response was almost indistinguishable (Figure 4B). When analyzed along the cell polarity axis (that is <FRET>front/rear; Supplemental Figure S2C), this negative response appeared somewhat more exaggerated in the cell front than the rear (Figure 4C). We observed no clear correlation between the FRET signal difference and cell velocity along the y-axis (Figure 4D). Remarkably, when the data were normalized by the basal activity that is average (<FRET> y+/y-) before stimulus ( t = -1 to 0 min), we discovered that the response between the [y+] and [y-] side completely overlapped with one another in both the timing and the amplitude (Supplemental Figure S5, A-B). This indicates that HL-60 cells exhibit a fold-change (Goentoro and Kirschner, 2009)in Cdc42 activity in response to a fast-wave stimulus. Along the cell polarity axis, the fold change showed a stronger undershoot at the rear of the cell in the descending phase (Supplemental Figure S5C), during which the absolute velocity decelerated (Supplemental Figure S5D). After wave passage, there was an increase in the cell velocity in the [y+] direction (Figure 4D; Supplemental Figure S5B; t > 1 min). To summarize the response to the 1 min wave stimulus, Cdc42 activity increased in the wavefront; however, there was no spatial asymmetry aligned with the axis of stimulus propagation. In other words, Cdc42 activity increased homogeneously over a pre-existing bias, which is consistent with the fact that cell orientation was unaltered (Figure 1O). Interestingly, during waveback, Cdc42 activity decreased more markedly on the front side of the cell (Supplemental Figure S5C), suggesting its possible role in cell deceleration.

**FIGURE 4:**
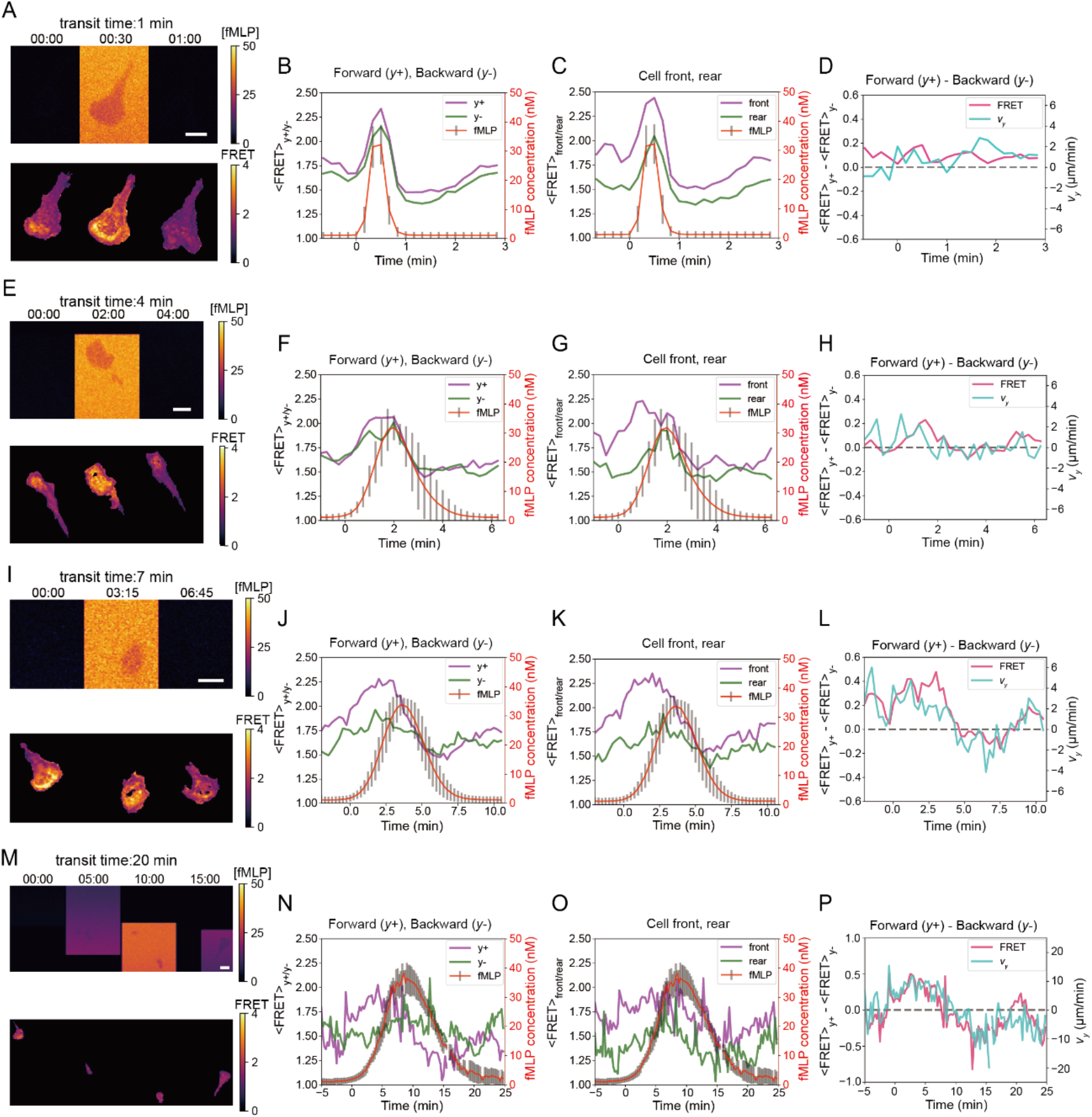
Spatio-temporal change in the Cdc42 activity is highly correlated with the wave speed dependency. FRET measurements under wave transit time of 1 min (A-D), 4 min (E-H), 7 min (I-L) and 20 min (M-P). (A-D) Snapshots of Rachu-Cdc42 FRET signal (lower panels) and the stimulus profile (upper panels) (A, E, I and M). Mean <FRET>y+ (magenta) and <FRET>y- (green) side along the y-axis (B, F, J and N). Mean <FRET>front (magenta) and <FRET>rear (green) taken along the cell velocity vector (see Supplemental Figure 2B, C for notation) (C, G, K and O). The mean of <FRET>y+ - <FRET>y- (magenta) plotted together with y-velocity (cyan) (D, H, L and P). n = 18 (B-D), 7 (F-H), 10 (J-L), and 13 cells (N-P).

Under the 4 min waves (Figure 4E), Cdc42 activity became more asymmetric compared to 1 min wave. The mean Cdc42 activity peaked at approximately 1 to 2 min on both [y+] and [y-] sides, which then exhibited a minor undershoot at 2 to 3 min before slowly returning to the pre-stimulus level (Figure 4F). In comparison, the difference in Cdc42 activity between the cell front and the rear was more pronounced (Figure 4G). The timing of the negative response coincided with a decrease in cell velocity (Figure 4H; see also Figure 1L). Under 7 min waves (Figure 4I), the spatial bias in Cdc42 activity became more aligned with stimulus orientation (Figure 4J), and the change in cell speed was more substantial (Figure 4K). Given that this matched well with the Cdc42 activity difference measured along the cell polarity axis (Figure 4K), the data indicated that cells were more aligned along the stimulus orientation in the 7 min wave (Figure 4, J and K) than in the 4 min wave (Figures 4, G and K). Accordingly, along the y-axis, the cell velocity also shows a good match with the difference in Cdc42 activity (Figure 4L). Given that the chemotaxis index in the wavefront of the 7 min wave was higher than that of the 4 min wave (Supplementary Figure S1, A and B), these features are in line with the notion that the Cdc42 activity gradient is pivotal in determining cell directionality.

The fold-change response (Supplemental Figure S5, I-L) similarly illustrated the features described above, which were based on the absolute mean Cdc42 activity (Figure 4J). A noticeable difference was that the average fold change was higher towards the [y-] side during the waveback (Supplementary Figure S5I). In contrast, there was almost no difference in the fold-change response when comparing the front and rear of the cell (Supplementary Figure S5K), indicating that the fold-change component of the Cdc42 activity gradient more directly reflects stimulus orientation rather than cell polarity. FRET measurements for the 20 min wave were difficult to conduct, as they required long-term tracking of cells at high magnification while simultaneously maintaining wave propagation in a consistent manner. Nevertheless, from a small number of samples, we were able to obtain patterns similar to those observed under the 7 min wave (Figure 4, M-P), except that the gradient of absolute Cdc42 activity between the [y+] and [y-] sides in the waveback could be detected (Figure 4N; 7–10 min), suggesting a good alignment between orientation as determined by the Cdc42 activity gradient and cell polarity. Accordingly, the fold-change pattern (Supplemental Figure S5, M, and N) was similar to the absolute Cdc42 activity readout (Figure 4, N, and P). These observations are consistent with the fact that the cells fully reverse-migrated under a 20 min wave. The fold change was higher in the cell rear than in the front (Supplemental Figure S5, O and P) because of our small sample set, which was skewed in the initial cell orientation.

### Cdc42 activation is sufficient to induce leading edge formation and its moderate suppression abolishes cell migration in the wavefront of fast waves

To further analyze the role of Cdc42, we adopted an optogenetic approach previously used in RAW.264 cells(Guntas *et al*., 2015). The system employs membrane translocation of the DH/PH domain of Cdc42GEF ITSN1, (O’Neill *et al*., 2016) which is known to interact with Cdc42(WT) and dominant-negative Cdc42(T17N), but not with inactive Cdc42(Q61L), Rac1, or RhoA(Hussain *et al*., 2001). We generated HL-60 cells stably expressing ITSN1(DH/PH) fused to tgRFPt-SspB (WT) and its binding partner, Venus-LOV-SsrA, which was tagged with a CAAX tail for plasma membrane anchoring (Figure 5A). When briefly stimulated uniformly with blue light, ITSN1(DH/PH)-tgRFPt-SspB localized to the plasma membrane within 6 s and then redistributed in the cytosol within 90 s (Supplemental Figure S6, A and B). Brief illumination of a small subcellular region elicited a spatially restricted membrane translocation of ITSN1(DH/PH)-tgRFPt-SspB (WT), followed by the formation of a membrane protrusion (Figure 5B). When iterated, local illumination elicited local membrane elongation and the cells were oriented accordingly (Supplemental Figure S6C). Local translocation of ITSN to the front and lateral sides (Figure 5C) as well as to the rear (Figure 5D) induced a new cell front. In addition, under a stationary gradient of fMLP (Figure 5E), a new cell front was induced at the site of local ITSN1 accumulation, thus overriding the chemotactic cell orientation. These results are qualitatively consistent with a detailed analysis based on light-induced GPCR activation reported earlier, (Bell *et al*., 2021; Hadjitheodorou *et al*., 2021) and more directly highlight the sufficiency of localized Cdc42GEF in inducing the cell front and directional movement of HL-60 cells.

**FIGURE 5:**
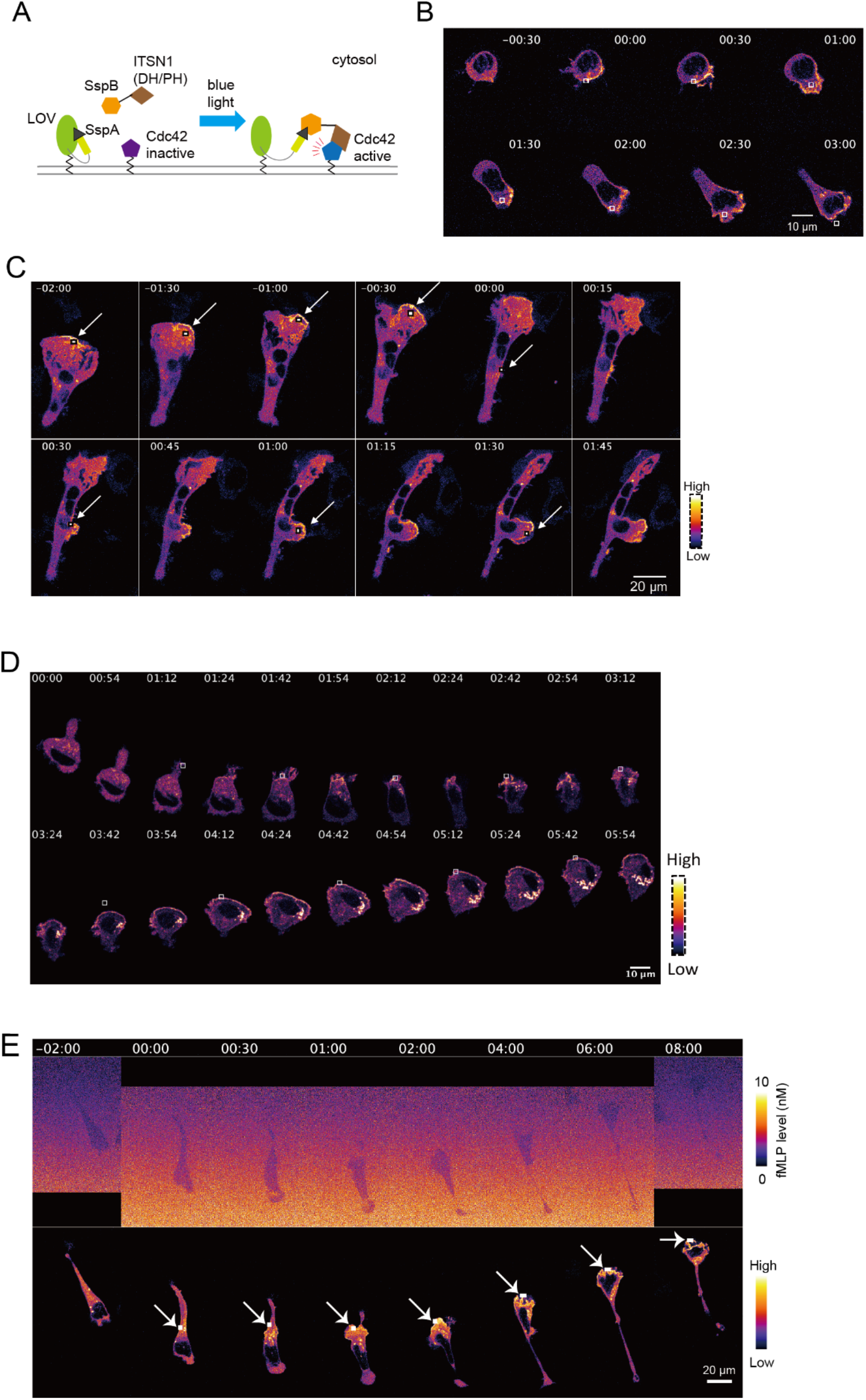
Local membrane accumulation of Cdc42GEF is sufficient for the leading edge formation. (A) Schematic representation of the iLID system. Dimerization of LOV-SspA and SspB-tgRFPt-ITSN(DH/PH) is induced by blue light. (B) Light-induced accumulation of SspB-tgRFPt-ITSN (DH/PH) and leading-edge formation in a non-polarized cell. (C) Light-induced accumulation of SspB-tgRFPt-ITSN(DH/PH) to the leading edge (t = -2 to 0 min) and at the lateral side (t = 0 to 1:30 min) promotes leading edge formation. (D) Light-induced accumulation of SspB-tgRFPt-ITSN(DH/PH) to the trailing end (t = 1:12 and 1:42) resulted in formation of a new leading edge. (E) Light-induced accumulation of SspB-tgRFPt- ITSN(DH/PH) in a chemotaxing cell (lower panel) in the presence of a fMLP gradient (upper panel). A 488 nm laser is applied to the white square region at 30 s intervals (B-E).

Next, we investigated the effect of Cdc42 inhibition on wave chemotaxis. Cdc42 knockdown causes severe defects in the establishment and maintenance of cell polarity(Bell *et al*., 2021). Accordingly, HL-60 cells treated with 20 µM ZCL278, a selective inhibitor of Cdc42 that targets Cdc42-ITSN(GEF) interaction(Friesland *et al*., 2013), completely abolished the migratory response to fast, intermediate and slow transit waves (Figure 6, A-C). Under a moderate treatment of 10 µM ZCL278, the overall chemotactic behavior was similar to that of non-treated cells (Figure 6, D-F) except that the mean velocity and the displacement were reduced markedly (Figure 6, G-L). At all transit times tested, the start of cell displacement in the wavefront was delayed and was not noticeable until the fMLP concentration reached approximately 10 nM. In addition, the response observed for the 7- and 20 min waves became more asymmetric (Figure 6, H and I), as was evident from the less sustained cell velocity at the waveback (Figure 6, K and L), thus resembling the response of non-treated cells under a 4 min wave (Figure 1H). In contrast, the cell velocity and its temporal profile showed little dependency on the wave speed (Figure 6, J-L). These data indicate that ZCL278-treatment has a more detrimental effect on the response to fast-wave stimuli (4 min transit time) than to slower waves (7 and 20 min), thus implicating the role of Cdc42 in response to relatively fast changes in attractant concentrations.

**FIGURE 6:**
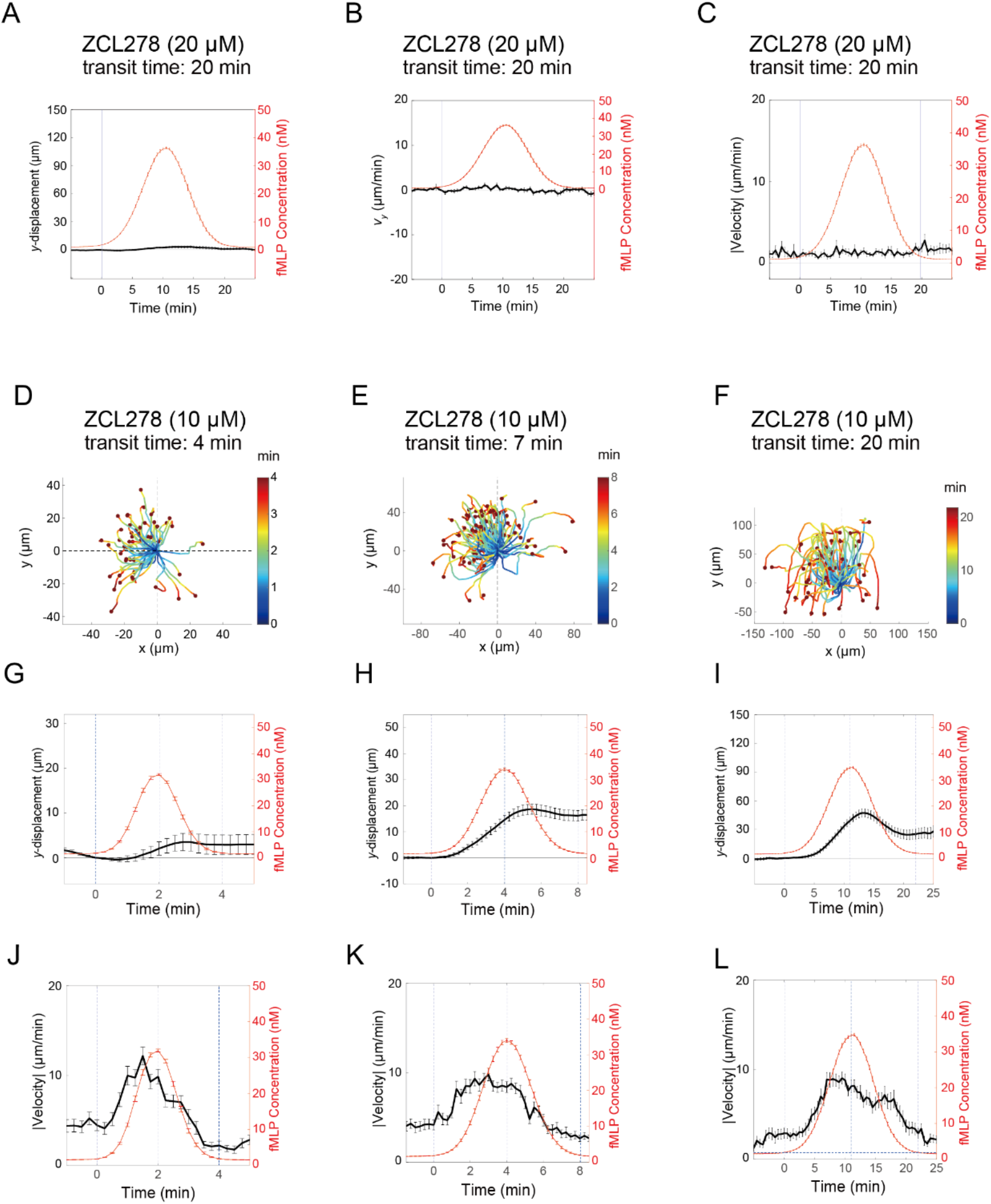
Moderate dosage of ZCL278-treatement affects response to the fast wave. (A-C) Strong ZCL278 treatment (20 µM). Average cell displacement in y-direction (A), centroid velocity (B), and absolute speed (C) in cells treated with 20 µM ZCL278 (n = 17). (D-L) Moderate ZCL278 treatment (10 µM). (D-F) Centroid trajectories. (G-I) Average cell displacement in y-direction. (J-L) Absolute speed. Wave transit time of 4 min (D, G and J), 7 min (E, H and K), and 20 min (F, I and L). n = 52 (G and J), 100 (H and K) and 60 cells (I and L).

### PI3K, Rac1, ROCK inhibition suppresses waveback response in slow waves

As moderate treatment with the Cdc42 inhibitor ZCL-278 strongly impaired cell migration under fast waves, but not slow waves, we hypothesized that a signaling module that acts in parallel with Cdc42 may play a role in a longer time-scale response. The GTPases Rac and PI3K are critical components for the formation of cell protrusions and cell polarity(Campa *et al*., 2015).

Optogenetic activation of PI3K induces cell protrusion and migration in a Cdc42-null strain, (Town and Weiner, 2023) thus hinting at an auxiliary layer of regulation. Elevated activity of Rac is known to occur after that of Cdc42 (Yang et al., 2015) and may constitute a positive feedback loop together with PI3K to enhance cell polarity(Srinivasan *et al*., 2003; Inoue and Meyer, 2008; Town and Weiner, 2023). Under the wave stimulus, The plekstrin homology domain of Akt (AktPH), which serves as a surrogate for the PI3K product PI(3,4,5)P3(Frech *et al*., 1997), exhibited a membrane translocation pattern that correlated well with cell directionality (Figure 7, A-C). AktPH-Clover translocated to the leading edge in the wavefront, which then became more uniform at the wave peak before becoming enriched in the new leading edge facing the reverse-oriented gradient (Figure, 7A and B; 7 min and 20 min wave), Figure 7C shows a case in which the cell did not show AktPH-Clover localization during the waveback and did not migrate backwards indicating a correlation between cell movement and the pattern of AktPH-Clover localization during fMLP wave stimulation. Figure 7, D-F show the average centroid displacement of cells treated with the PI3K inhibitor LY294002. Interestingly, while there was no noticeable effect under the 4 and 7 min waves (Figure 7, G and H), migration was strongly diminished in the back of the 20 min wave (Figure 7I). The changes in the average cell speed appeared to be similar across all wave speeds, that is, the velocity profile peaks at the midpoint of the wavefront (Figure 7, J-L). This contrasted with the prolonged increase in cell speed in untreated cells (Figure 1, M and N). Dependency on the pre-stimulus orientation (Figure 7, M-O) was similar to that of the non-treated cells (Figure 1, P-R), except that backward-moving cells were better at showing positive displacement in the wavefront (Figure 7, M and N; blue), indicating that LY294002 helps cells override past movement. These features were also observed in cells treated with the Rac1 inhibitor NSC23766 (Supplemental Figure S7), suggesting that Rac1 and PI3K play similar roles in sustaining cell velocity at the back of slow waves.

**FIGURE 7:**
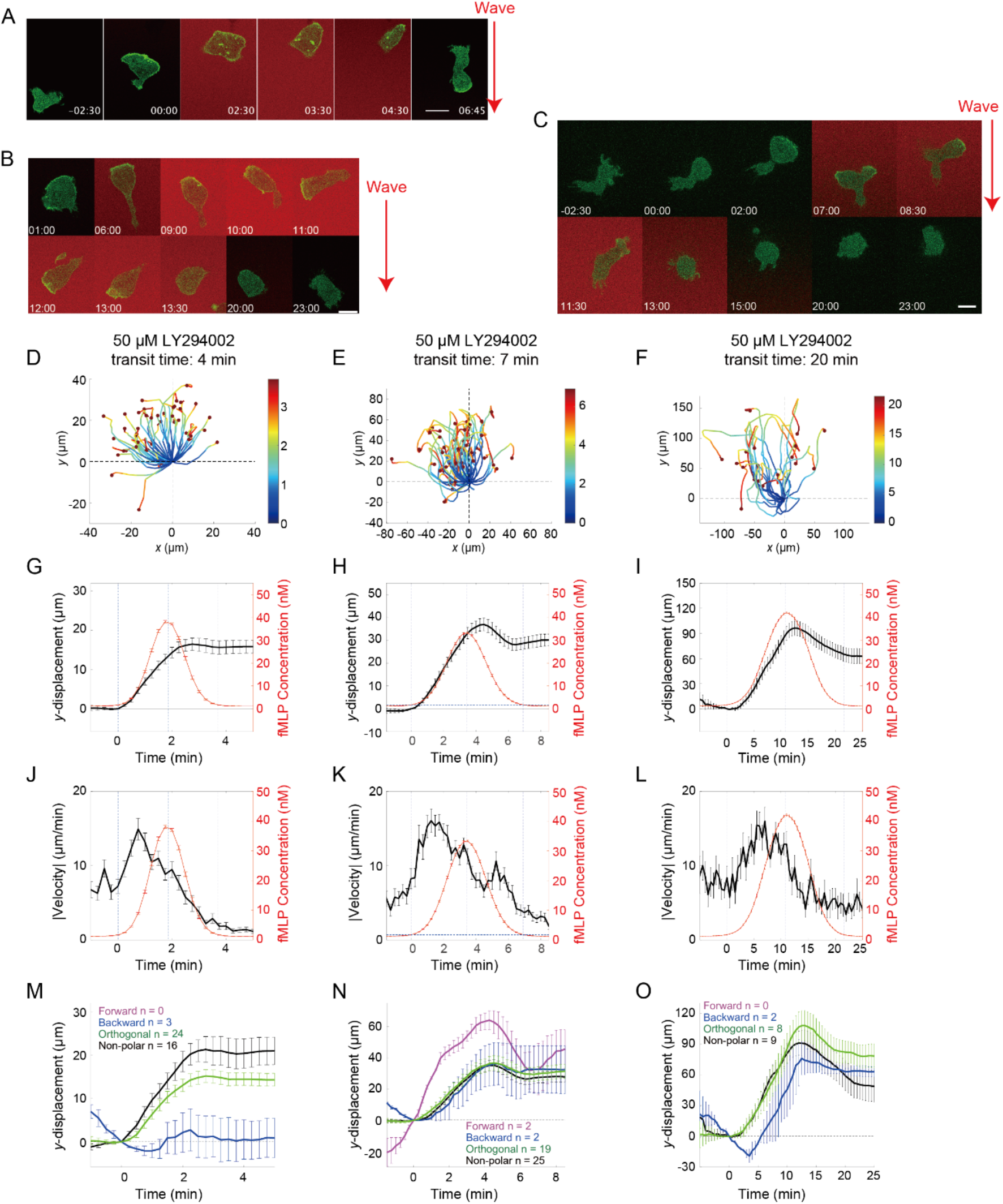
LY204002-treated cells exhibit poor re-orientation in the slow wave. (A-C) Membrane translocation of PHAkt-Clover under the fMLP wave stimulus. Wave transit time: 7 min (A; representative data out of n = 10 cells) and 20 min (B and C; representative data out of n = 5 cells). (D-F) Centroid trajectories. (G-I) Average cell displacement in y-direction. (J-L) Absolute speed. Wave transit time of 4 min (D, G and J), 7 min (E, H and K), and 20 min (F, I and L). n= 43 (G and J), 48 (H and K) and 19 cells (I and L). (M-O) The mean y- displacement of cell centroids based on the initial cell orientation for wave transit time of 4 min (M), 7 min (N) and 20 min (O). Forward (magenta), Backward (blue), Orthogonal (green) and Non-polar (black); see Supplemental Figure S2 for notation.

These results are in stark contrast to what was observed for ZCL278-treated cells, where the migratory pattern was disrupted only at fast to moderate wave speeds (therefore cell migration in the wavefront). Moderate ZCL278 treatment deteriorated cell orientation as measured by the chemotaxis index in the wavefront, especially for the fast wave (Supplemental Figure S8, A-C), while the average cell velocity was not strongly affected (Figure 6, J-L). In contrast, treatment with either LY294002 or NSC23766 did not affect cell orientation, as judged by the chemotactic index (Supplemental Figure S8, D-I). If we plot Cdc42 activity difference between the [y+] and [y-] side (<FRET> y+ - <FRET> y-) against the cell speed in the y-direction (Figure 8, A-D), there was a high correlation between the two before and after the wave stimulus (Figures 8, A and D; correlation coefficient 0.73 and 0.66). This suggests that intracellular gradient of Cdc42 activity is highly correlated with the orientation of random cell movements. However, the correlation was substantially weaker under the wave stimulus (Figure 8, B and C) (correlation coefficients 0.43 and 0.39). The Cdc42 gradient appears to lose correlation with cell velocity in an arbitrary direction as the overall trend was also seen along the x-axis (Figure 8, E-H correlation coefficients 0.59, 0.49, 0.47, and 0.68); see Supplemental Figure S2D for notation <FRET>x+/x-). This appears to be accompanied by high velocity points in 20 min wave data (Figurel 8B and C; red) which were also present after stimulation (Figure 8D; red) suggesting that a build-up of memory further reduces the correlation between cell velocity and the Cdc42 activity gradient under slow waves.

**FIGURE 8:**
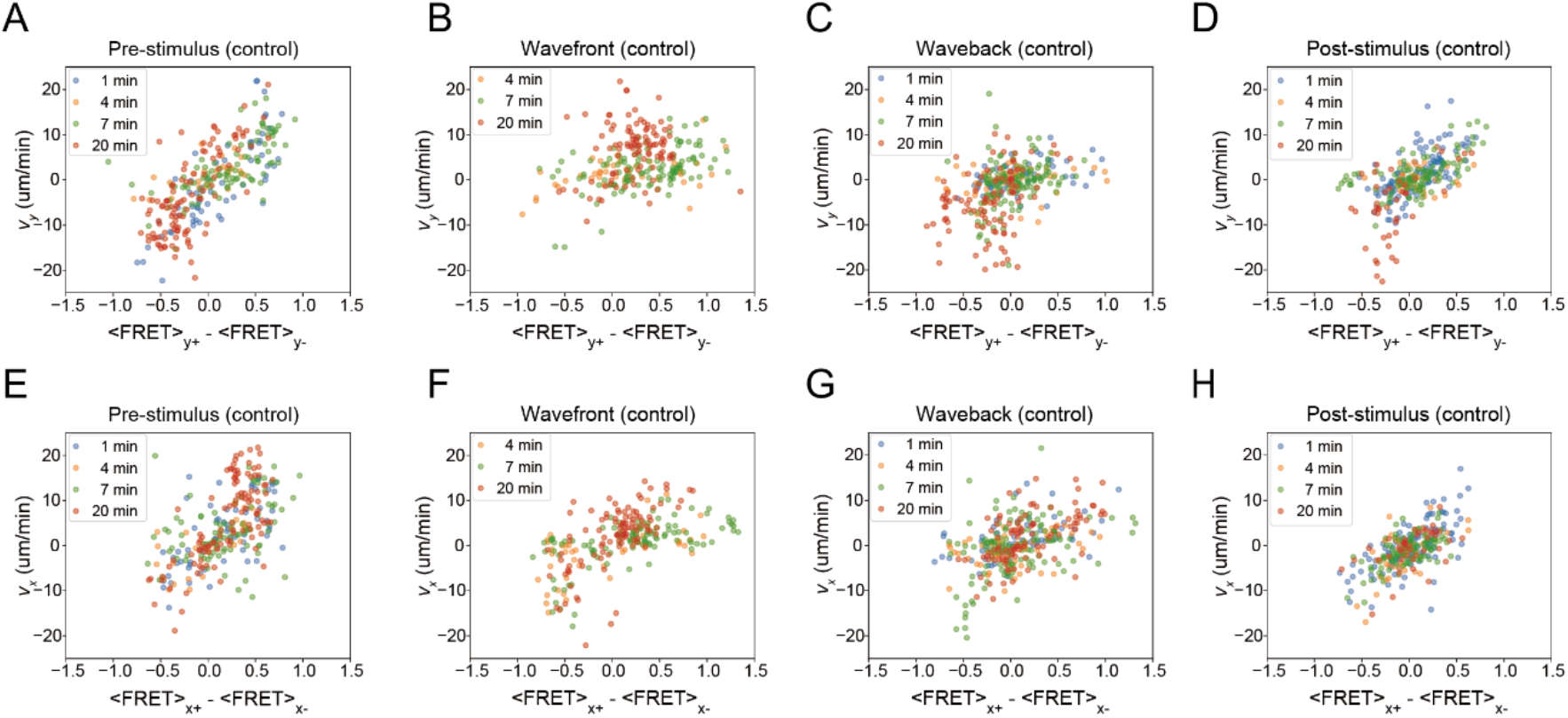
Correlation between centroid speed and Cdc42 activity. (A-D) Velocity along the y-axis in the wave stimulus without inhibitors is plotted against the Cdc42 activity difference between the [y+]-side and [y-] side at the time frames (A) before stimulation, (B) in the front side of the wave, (C) in the back side of the wave, and (D) after stimulation. (E-H) Velocity along the x-axis in the wave stimulus without inhibitors is plotted against the Cdc42 activity difference between the (x+)-side and (x-) side at the time frames (E) before stimulation, (F) in the front side of the wave, (G) in the back side of the wave, and (H) after stimulation. Colors indicate different wave transit time. Data samples are the same as Figure 4.

To further address cell re-orientation or its lack in the waveback, we analyzed cells treated with an inhibitor of Rho-GTPase-associated protein kinase (ROCK), which plays a major role in regulating interactions between actin and myosin through the inhibition of myosin light chain phosphorylation(Niggli, 1999; Xu *et al*., 2003; Hadjitheodorou *et al*., 2021). We found that cells treated with the ROCK inhibitor Y-27632 (Uehata *et al*., 1997) also failed to reverse-migrate during the waveback (Figure 9, A-H, M-P). For those that made a turn, U-turns and polarity reversals showed similar occurrences (Supplemental Figure S3). The average speed at waveback also decreased markedly (Figure 9, I-L). This was surprising, as ROCK inhibition should help sustain myosin II accumulation at the cell rear and thus facilitate polarized cell movement, as demonstrated in 1D channels(Prentice-Mott *et al*., 2016; Hadjitheodorou *et al*., 2021). As the trend continued even under a slower 30 min wave stimulus, the effect was likely not due to a delay in the timing of gradient reversal. Owing to the large drop in cell velocity, the chemotaxis index in the waveback showed a median value close to zero (Supplemental Figure S8, J-L; Ttransit/4), which was distinct from the LY294002- or NSC23766 treated cells. Cdc42 activity in Y-27632 treated cells was suppressed at the back of the 20 min wave (Figure 10A), and thus, the overall time development closely resembled that of non-treated cells under 4 to 7 min wave stimulus (Figure 4, F and J). This suggests the importance of ROCK in sustaining Cdc42 activity at the back of the slow waves. When we compared along the stimulus axis the cell centroid velocity (Figure 10B) and the difference in Cdc42 activity (Figure 10, C-F), the correlation in the waveback increased in Y-27632-treated cells (correlation coefficient 0.71; Figure 10E) compared to that in non-treated cells (Figure 8C), suggesting that there is a basal velocity component that is Cdc42-dependent but ROCK-independent, and that this is masked by ROCK-dependent regulation under prolonged stimulation.

**FIGURE 9:**
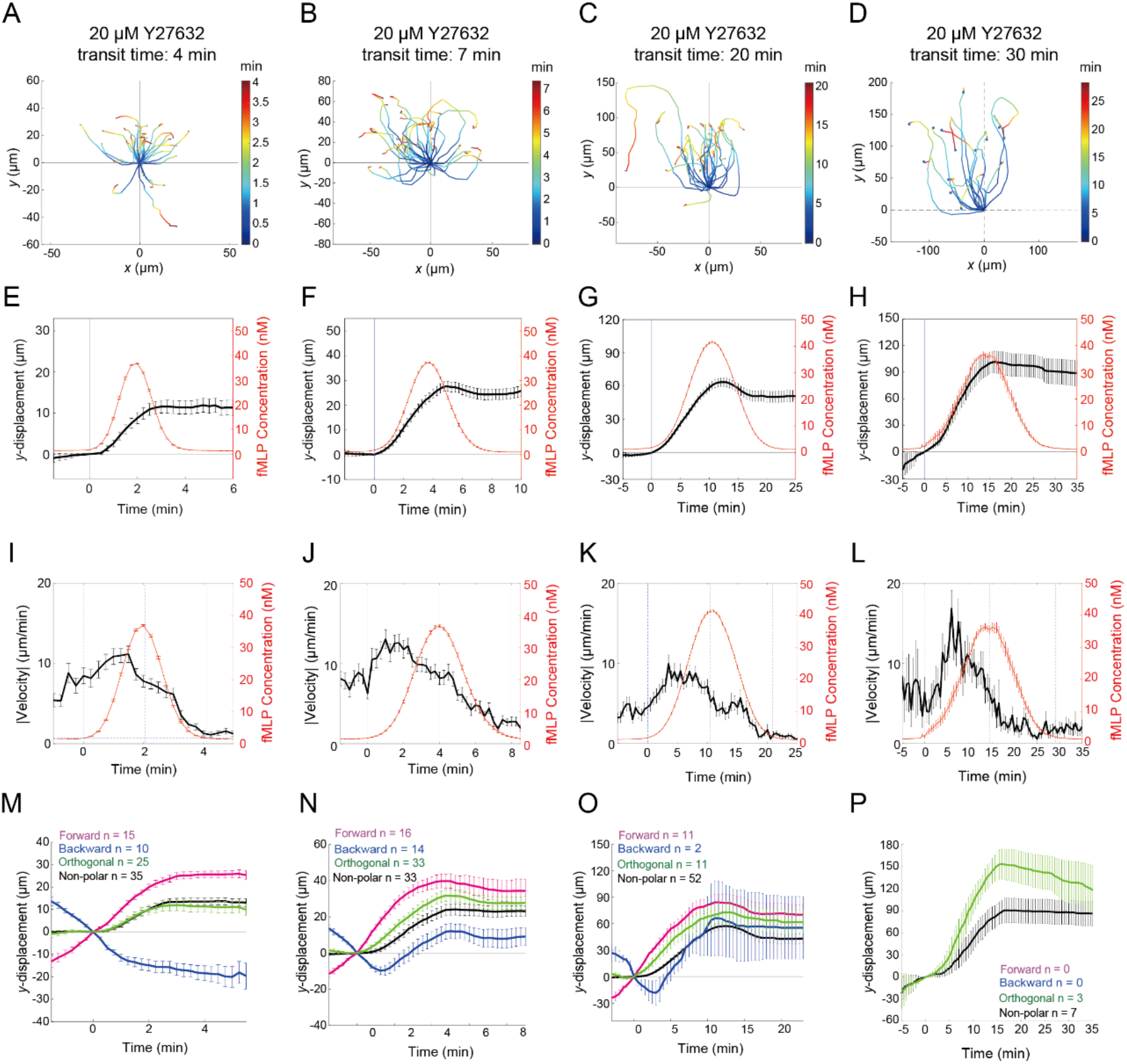
Y-27632-treated cells exhibit poor re-orientation in the slow wave. (A-D) Centroid trajectories. (E-H) Average cell displacement in y-direction. (I-K) Absolute speed. Wave transit time of 4 min (E and I; n = 89 cells), 7 min (F and J; n = 96 cells), 20 min (G and K; n = 76 cells), and 30 min (H; n = 15 cells). (M-P) The mean y-displacement of cell centroids based on the initial cell orientation for wave transit time of 4 min (M), 7 min (N), 20 min (O), and 30 min (P). Forward (magenta), Backward (blue), Orthogonal (green), and Non-polar (black); see Supplemental Figure S2A for notation.

**FIGURE 10:**
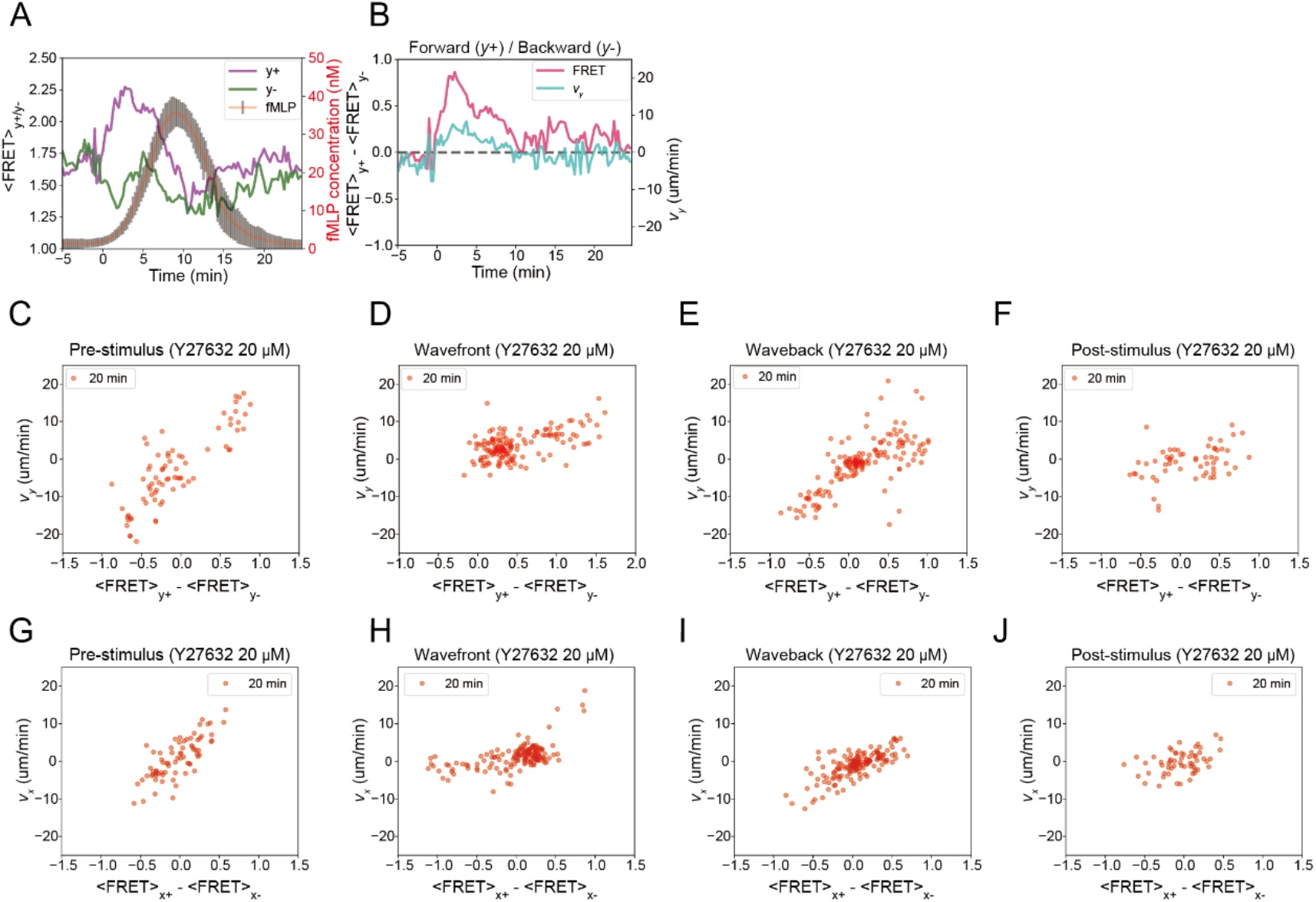
Correlation between centroid speed and Cdc42 activity is not interrupted by the 20-min wave stimulus in Y-27632 treated cells (n =4 cells). (A) Cdc42 activity difference between the (+)-side and (-) side along the y-axis during 20 min transit wave. (B) Time development of centroid velocity and Cdc42 activity difference between the [y+]-side and [y-] side. (C-F) Velocity along the y-axis in the wave stimulus in Y-27632 treated cells is plotted against the Cdc42 activity difference between the [y+]-side and [y-] side at the time frames (C) before stimulation, (D) in the front side of the wave, (E) in the back side of the wave, and (F) after stimulation. (G-J) Velocity along the y-axis in the wave stimulus in Y-27632 treated cells was plotted against the Cdc42 activity difference between the [x+]-side and [x-] side before stimulation (G), during the wavefront (H), the waveback (I), and after stimulation (J).

## DISCUSSION

The present study demonstrated that HL-60 cells on average are capable of migrating unidirectionally in the direction opposite to that of attractant wave propagation (Figure 11). For both fMLP and LTB4, uni-directional migration occurred at intermediate speeds of approximately 100 to 170 µm/min. When the wave was slower, the HL-60 cells re-oriented and chased the passing wave; therefore, the net displacement was almost zero or even negative. Under fast waves, the stimulus can reset pre-existing polarity, but the cells cannot define a new directionality. These properties reveal the time-scale dependence of elementary input-output relationships that underlie HL-60 chemotaxis. If the attractant rises quickly, a transient rise in Cdc42 occurs which attenuates along with the cell velocity well before the stimulus peaks. If the attractant concentration increases slowly, the cell speed remains elevated, which depends on PI3K/Rac and ROCK. As for waveback, if the attractant descends quickly enough, cell orientation is randomized and cell speed is reduced (except for the LTB4 inverted wave), which is not dependent on PI3K/Rac or ROCK, but rather on the temporal response of Cdc42. If the rate of attractant decrease is slow, Cdc42 gradient persists which depends on ROCK.

**FIGURE 11:**
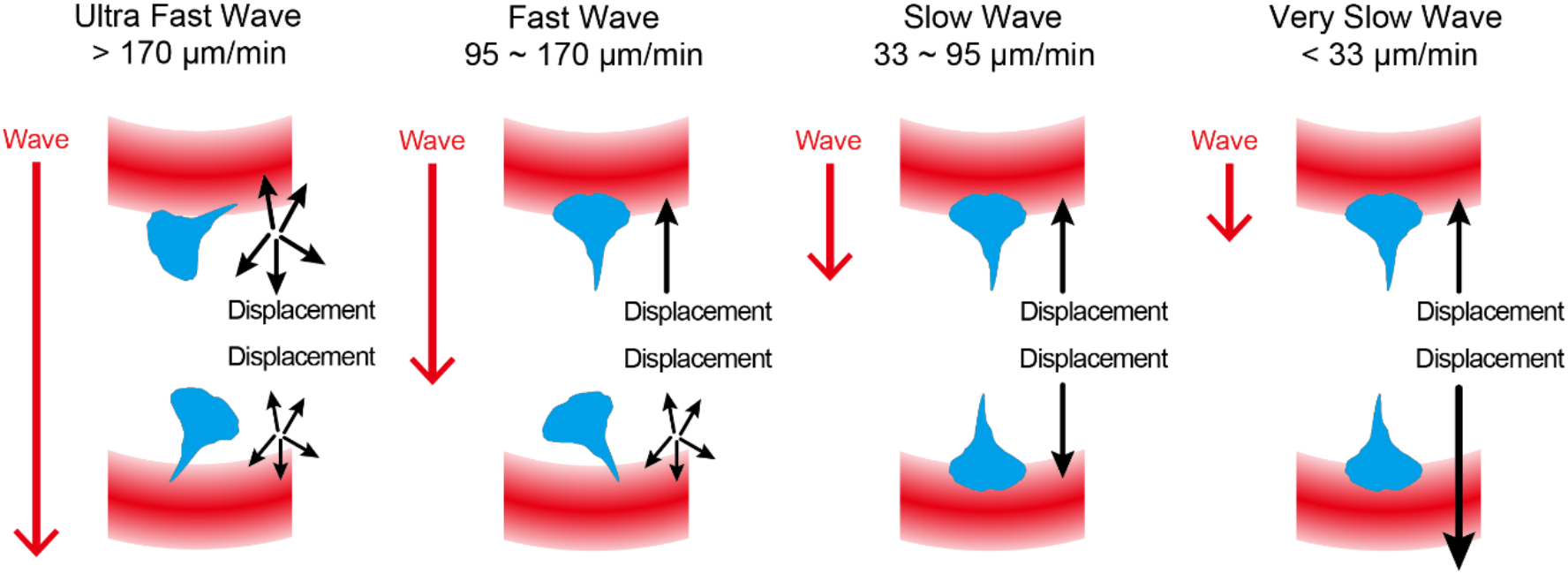
Summary of the wave-speed dependency in the migratory response. Red arrows indicate the direction of wave propagation. Black arrows indicate cell orientation and the magnitude of displacement. Response to a wavefront and a waveback are shown at the top and the bottom, respectively.

Although the present study does not provide experimental support to decipher the origins of speed dependence, several features are worth discussing in light of the candidate scenarios. In the 4- and 7 min waves, Cdc42 activity was biased along the wave stimulus axis (Figure 4, F and J). The bias was less clear at 4 min and deviated only slightly from the front-back bias (Figure 4, G and K), indicating that the response was more spatially uniform compared to the 7 min wave. This agrees with the fact that backward-moving cells in the wavefront of the 4 min wave stopped but were not able to re-orient (Figure 1P). If one assumes a first-hit scheme that realizes a one-side wins all-type response, our observation hints at the speed of inhibition required to spread within a single cell. If we assume that activation step is local, speed of fast wave (transit time of 1 to 2 min) close to or above 300 to 600 µm/min is interpreted as having won the race of stimulating the other sides of a cell before the long-range inhibition kicked in. If this long-distance effect was mediated by diffusive signal that emanated from the side first hit by the attractant, a calculation based on the mean square displacement within a 2D sphere predicts a diffusion constant of roughly 15 to 30 µm^2^/s which is typical for a cytosolic protein(Nakajima *et al*., 2014). However, this estimate does not rule out tension as being the inhibitory signal, as the speed of the fast wave translates to approximately 1–2 s per cell diameter, which is also comparable to the time of the spread of tension, which takes approximately 1–10 s(De Belly *et al*., 2023).

According to the local excitation global inhibition (LEGI) model(Iglesias and Levchenko, 2002; Takeda *et al*., 2012; Nakajima *et al*., 2014; O’Neill and Gautam, 2014; Skoge *et al*., 2014), localized activation and long-range inhibition are assumed to occur according to the degree of receptor-attractant binding in a feed-forward manner. Owing to long-range inhibition, the transient response in the LEGI model can support a first-hit response (perception of the time difference of stimulus arrival)(Nakajima *et al*., 2014). Note that this scheme can also convey a negative signal, that is, a temporal decrease in the mean attractant concentration. Under the LEGI framework, this would result in local “de-activation” and global “de-inhibition” and thus should also re-orient a cell. Our observations suggest that, although there is a small negative temporal response, it is not spatially biased along the stimulus axis (Figure 4J). This contrasts with the almost complete absence of an undershoot response for Ras in *Dictyostelum.* In contrast, in slow waves, the cells re-orient in the waveback. This can be interpreted as a stationary-state response in the LEGI framework, where the leading-edge response is proportional to the relative receptor occupancy. According to this framework, the results of pharmacological perturbations suggest that the Cdc42 gradient in this near-stationary state is not substantially amplified and is too shallow to drive re-orientation.

An earlier optogenetic study showed that the front-to-back gradient of Cdc42 and Rac is generated by local activation and global inhibition; however, this inhibitory signal may not need to diffuse globally from a localized source(de Beco *et al*., 2018). If we do not assume stimulus-induced long-range inhibition, sensing the spatial gradient (that is, steepness) in the strict sense should be difficult because the signaling network is not provided with information regarding the spatial average. Alternatively, signal perception can be purely temporal in origin(Vicker *et al*., 1986). One interesting possibility is that spatial gradient information can be decoded based on space-time translation (Gerisch *et al*., 1975) using a “pilot pseudopods”(Devreotes and Zigmond, 1988; Town and Weiner, 2023). This mechanism can, in principle, explain why cells did not re-orient in the wave back, as no matter which direction the pilot pseudopods are extended, they should all perceive a temporally negative signal. As the wave speed decreased, the pilot pseudopod began to win the speed competition and should be able to detect temporally positive signals along a near-stationary gradient. However, from our wave chemotaxis results, it requires pilot pseudopod to extend at least at the speed equivalent to the 20 min wave (that is, 33 µm/min) which exceeds the speed of filopodia (∼1 µm/min) (Mogilner and Rubinstein, 2005) and pseudopod extension (10 µm/min) (Fritz-Laylin *et al*., 2017). At least in *Dictyostelium*, pilot pseudopods are not necessary, as latrunculin-treatment did not affect directional Ras activation under traveling-wave stimuli(Nakajima *et al*., 2014). Similar conditions were difficult to test for Hl-60 because of cell detachment under our flow conditions.

Another plausible explanation for the wave speed dependency is that there is an intrinsic time difference between the time required to form a new leading edge versus that to re-orient the pre-existing polarity, as the latter involves more time to re-construct cytoskeletal filaments. Optogenetic application of localized GEF induced a new leading-edge on all sides of the cell (Figure 5), suggesting that if there was long-range inhibition, our optogenetic manipulation was somehow able to escape or compete with existing inhibition. A reaction-diffusion scheme in which long-range inhibition is regulated downstream of the local activator to form a negative feedback loop that acts long range is more compatible with this observation. This scheme coupled with some form of positive feedback is compatible with the spontaneous breaking pattern of Cdc42 activity observed in randomly migrating cells(Yang *et al*., 2016). The first-hit scheme also applies in this scenario, because the initial stimulus position determines the position of the peak of the pattern. Here the unidirectional migration is interpreted as a ‘front-locking’ behavior (Meinhardt, 1999; Town and Weiner, 2023) predicted in such reaction-diffusion systems. In slow waves, one can envisage a local inhibitory signal that unlocks the leading-edge stability, as suggested(Town and Weiner, 2023). Further analyses are necessary to elucidate these possibilities.

Notably, the speed dependence observed in this study (Figure 11) was remarkably similar to that of *Dictyostelium* chemotaxis in response to a traveling wave of cAMP(Nakajima *et al*., 2014). This was surprising given that the common ancestors of metazoans and amoebozoa diverged more than 1 billion years ago(Parfrey *et al*., 2011; Fiz-Palacios *et al*., 2013). Similarities in chemotaxis between the two systems must entail constraints on signaling pathways and cytoskeletal architectures from common ancestry and convergent traits that evolved independently(Artemenko *et al*., 2014; Filić *et al*., 2021). The former can be recognized from large overlaps in actin-related proteins and signaling networks. They are both G-protein coupled receptor (GPCR)-mediated and exhibit adaptive responses in small GTPase and PI3K activity, (Artemenko *et al*., 2014; Tang *et al*., 2014) in addition to excitable dynamics that drive spontaneous membrane protrusions(Tang *et al*., 2014; Yang *et al*., 2016). These signaling characteristics can be considered breeding grounds for reaction-diffusion dynamics, which can be fitted to a plethora of response characteristics in a context-dependent manner. In contrast to the many similarities found in the regulation of leading-edge formation, the two systems may diverge in terms of how the trailing edge is defined (Filić *et al*., 2021). In neutrophils, RhoA and ERM are key components that regulate the trailing tail (Hind *et al*., 2016; Prentice-Mott *et al*., 2016). Although *Dictyostelium* possesses a distant RhoA homologue, RacE, there is no ROCK or ERM protein, and there is no evidence that microtubules are involved in the maintenance of cell polarity. Nevertheless, myosin regulation is pivotal for wave chemotaxis in *Dictyostelium* as is evident from the fact that the loss of the myosin heavy kinases MhckA or MhckC results in cells that cannot re-orient in the wavefront(Wessels *et al*., 2012). Myosin heavychain kinases act to disassemble myosin filaments (Yumura *et al*., 2005) and a recent study (Kuhn *et al*., 2024) reported using rapamycin induced dimerization, that MhckC inhibits Ras and PI3K activity thus implicating an actomyosin-dependent positive feedback loop.

Considering the biological implications of the present results, it is tempting to speculate that LTB4 and other attractants travel in pulses *in vivo* to guide neutrophil swarming. The lessons learned from *Dictyostelium* suggest that a few conditions must be met for traveling pulses to occur(Goldbeter, 1996; Gregor *et al*., 2010). 1) LTB4 binding to the membrane-bound receptor accelerates the release of LTB4; 2) the response must adapt under prolonged receptor-ligand binding, thus stopping LTB4 release; 3) extracellular LTB4 is degraded and cleared; and 4) the system ‘de-adapts’ and becomes responsive again. The autocrine stimulatory loop of LTB4 is known to be mediated by arachidonic acid in human neutrophils, and this can be blocked by an LTB4 receptor antagonist, as well as LY223982(Surette *et al*., 1999). Also, in human neutrophils, extracellular LTB4 is known to be rapidly degraded by ω-oxidation making its half-life short; from seconds to minutes (Archambault *et al*., 2019) close to the time-scale of cAMP hydrolysis by phosphodiesterase in *Dictyostelium* which is essential for cAMP oscillations(Gregor *et al*., 2010). As for response adaptation, the activation of Cdc42 subsided under persistent stimulation by LTB4, suggesting an adaptation mechanism at or near the receptor. Adaptation may also be mediated by accumulation of LTB4 metabolites, which downregulate LTB4-mediated responses(Archambault *et al*., 2019). Owing to the cyclic nature of steps 1) through 4), a site that gives rise to a pulse first is likely to give rise to another pulse; thus, periodic pulses can self-organize. A recent in vitro study on human neutrophils demonstrated the occurrence of repeated pulses of traveling Ca^2+^ waves, which were highly correlated with the direction and timing of cell movement(Strickland *et al*., 2023). As an increase in cytosolic Ca^2+^ promotes LTB4 production, Ca^2+^ waves may reflect the timing of LTB4 release. There, the speed of Ca^2+^ wave that accompanied cell aggregation was 60–120 µm/min (Strickland *et al*., 2023) which is in good agreement with the wave speed that supported uni-directional migration in our present study. According to Fisher’s equation, the wave speed is a function of the diffusion constant and rate of excitation. By reducing the rate of LTB4 release, the speed of wave propagation should start to slow, and aggregated cells should start to show bidirectional migration. As the waves slow down further, the cells continue to chase the back of the wave and thus disperse (Figure 11). A similar directional change, compatible with these movement patterns, has been noted for neutrophil chemotaxis during the resolution of inflammation in zebrafish(Mathias *et al*., 2006; Deng and Huttenlocher, 2012).

## Materials and Methods

### Plasmid and transformation

An expression vector for Akt-PH-Clover was constructed as follows: The DNA encoding Clover was PCR-amplified using pcDNA3 Clover (Lam *et al*., 2012)(Addgene #40259; a gift from Professor Michael Lin at Stanford University) as a template. GFP in the pcDNA3 Akt-PH-GFP plasmid (Kwon *et al*., 2007)(Addgene #18836; a gift from Professor Craig Montell at UC Santa Barbara) was replaced with Clover using the HindIII/BamHI site. To obtain the Akt-PH-Clover expression plasmid, Akt-PH-Clover was PCR-amplified from pcDNA3 Akt-PH-Clover and subcloned into the Epstein-Barr virus replicon-based plasmid pEBMulti Neo (WAKO 057- 08131). To obtain HL-60 cells stably expressing Akt-PH-Clover, pEBMulti Neo Akt-PH-Clover was introduced into the cells using a 2-step pulse electroporator (NEPA21; Nepa Gene). G418 (Invitrogen 11811023) was added to the medium at a final concentration of 1 mg/mL.

The piggyBac vector pPBbsr containing RaichCdc42 (pPBbsrRaichuCdc42) (Komatsu *et al*., 2011; Aoki *et al*., 2012) was a gift from Professors Kazuhiro Aoki and Michiyuki Matsuda at Kyoto University. For the optogenetic control of Cdc42, Venus-iLID, and ITSN1-tgRFPt-SspB were PCR-amplified from pLL7.0:Venus-iLID-CAAX (Addgene #60411; a gift from Prof. Brian Kuhlman at the University of North Carolina at Chapel Hill) and pLL7.0:hITSN1(1159-1509)- tgRFPt-SspB (Addgene #60419 a gift from Prof. Brian Kuhlman), respectively(Guntas *et al*., 2015). The PCR products were ligated into the pPBbsr. To obtain stable cell lines, the piggyBac expression vector and Super PiggyBac transposase expression vector (PB210PA-1, System Bioscience) were introduced into the cells using a 2-step pulse electroporator (NEPA21; Nepa Gene, Ltd.). Blasticidin S (R210-01; Invitrogen) was added to the medium at a final concentration of 5 µg/mL for selection.

#### Cell culture and preparation

HL-60 cells (RCB 0041; RIKEN BRC) were cultured at 37 °C and under 5% CO2 in an RPMI- 1640 medium containing L-glutamine, phenol red, and 25 mM HEPES (Wako 189-02145) supplemented with 10% heat-inactivated fetal bovine serum (Sigma-Aldrich 172012) and an antibiotic–antimycotic mix (Sigma-Aldrich A5955). The cells were diluted in fresh medium every 1–4 days and maintained below 1 × 10^6^ cells/mL. For cell differentiation, 1.3% dimethyl sulfoxide (Sigma-Aldrich D2650) was added to the medium, and the cells were cultured for 3–4 days.

#### Traveling wave stimulation

A commercially available flow chamber with 3 inlets and 1 outlet (µslide 3 in 1, Ibidi # 80311) was loaded with 150 µg/mL fibronectin (Corning 354008) solution in phosphate buffered saline (PBS, pH 7.4) and allowed to equilibrate for 1–3 h at 22–25 °C. The chamber was then removed from the coating solution and washed with PBS prior to use. Stock solution of 10 mM fMLP (Sigma F3506) and 1 mM LTB4 (Cayman 20110) were prepared in DMSO. To obtain a 1mM LTB4 stock solution, the LTB4 ethanol solution was purged with nitrogen gas to allow ethanol evaporation, after which LTB was immediately dissolved in DMSO. Differentiated HL-60 cells were collected and resuspended at a density of 1–3 x 10^5^ cells/mL in Hanks’ balanced salt solution HBSS(+) (WAKO 082-09365) containing 1 nM fMLP for the fMLP wave experiments. For LTB4 wave experiments, cells were suspended in plain buffer. For the inverted-wave experiments, the cells were suspended in 50 nM fMLP and 2 nM LTB4. The cell suspension was loaded into the chamber and the cells were allowed to attach to the substrate before time-lapse image acquisition. Wave stimulation was performed as described previously(Nakajima and Sawai, 2016). Specifically, a flow of buffer with peak and basal concentrations of the chemoattractant was provided from the center and side channels, respectively, using a programmable pressure regulator (MFCS, Fluigen) or syringe pumps (Pump 11 Elite, Harvard Apparatus). Source solutions of fMLP and LTB4 contained 0.4 µg/mL Alexa Fluor 594 (Invitrogen A10438, M.W. 758.8). See Figure 1A for the flow parameters. For consistency, time-lapse imaging was performed at a fixed region (0.32 mm wide and 2 mm long) positioned between 2 and 4 mm downstream from the center of the junction.

### Microscopy

An inverted microscope (IX-81; Olympus) equipped with a confocal multibeam scanning unit (CSU-X1; Yokogawa) and an EM-CCD camera (Evolve 512; Photometrics) were used for all live-cell imaging, except for those that involved optogenetic manipulation. For wave chemotaxis experiments using HL-60 cells, images were acquired every 15 s (except for the 1 min wave, which was at 10 s intervals) with a 10x objective lens (UPlanSApo NA 0.4, Olympus) by employing a 561 nm laser (25 mW Melles Griot) as a light source, a dichroic mirror (DM405/488/561), and a 575 nm long pass filter (BA575IF, Olympus) to visualize the stimulus delivery. Transmitted light images were obtained sequentially through the bright-field light path in the scanning unit, using a tungsten halogen lamp for illumination. In addition to the red fluorescence channel and the transmitted light images, for PHAkt-Clover/HL-60 cells, green fluorescence images were obtained with a 60 x objective lens (UPlanSApo NA 1.35) using 488 nm laser (50 mW, Vortran Laser Technology) and a 510-550 nm bandpass filter (BA510-550).

CFP and YFP channel images were obtained using a 445 nm laser (40 mW, Vortran Laser Technology) with a 60 × objective lens, a triple-band dichroic mirror (DM445/515/561), and an emission filter (FF01-542/27-25 Semrock). Confocal images were focused at around z = 3–5 µm from the substrate surface. All experimental conditions presented in this study were repeated at least two to three times.

For optogenetics using local blue-light illumination and simultaneous fluorescence imaging, an inverted microscope (IX-83; Olympus) equipped with a confocal multibeam scanning unit (CSU-W1; Yokogawa), an EM-CCD camera (iXon Ultra888; Andor) and a photoactivation unit (FRAPPA, Andor) was used. Time-lapse images were acquired at 15 s intervals (except for Supplemental Figure S6, A and B, which were acquired at 6 s) with a 60 × objective lens. For imaging, a 561 nm laser (100 mW); and 638 nm laser (140 mW, Cobolt) were used as light sources in conjunction with a dichroic mirror (DM405/488/561/640) and emission filters (FF02-617/73- 25, FF01-685/40-25 Semrock). A 488 nm laser (100 mW) was used as the light source for blue light illumination. Using the photoactivation unit, blue-light was applied at 30 s intervals to a rectangular region (20 × 20 pixels, 1 pixel = 0.22 µm) three times with 20 µs pulse duration per pixel.

#### Cell tracking and stimulus estimate

For cell tracking, cells that appeared severely attached to the substrate and could not migrate were excluded from the analysis. The trajectories of cells in the grayscale transmitted light images were manually tracked using the ImageJ plugin MTrackJ, (Meijering *et al*., 2012) except for high-magnification images (FRET data), which were automatically tracked using the ImageJ plugin TrackMate. The velocity was calculated as the change in the cellular position between neighboring time frames. The cell displacement was calculated by subtracting the y-axis coordinates of the cell centroids at the time of stimulus arrival (t = 0) from the y-axis coordinates at a given time point. The patterns of re-orientation were determined by manual inspection. The chemotactic indices of individual cell trajectories were quantified by taking the ratio of the signed y-displacement to the length of the 2D displacement at a time interval of 1 min. Patterns of pre-stimulus cell movement (‘backward,’ ‘forward,’ ‘orthogonal,’ and ‘non-polar’; Supplemental Figure S2A) were determined based on the centroid displacement between t = -1 to 0 min. Cells with displacement smaller than 5 µm was determined as ‘non-polar.’

‘Normalization image’ and ‘background image’ were captured at the observation region in a chamber filled with HBSS+ with or without 0.4 µg/mL Alexa Fluor 594, respectively. Brightness unevenness that derives from the optics and the camera was removed from the timelapse images (red-channel) by dividing background subtracting images by background subtracted ‘normalization image’.

The stimulus profile was obtained by spatially averaging a 101 × 3 pixel region surrounding the initial centroid of a cell. Estimates of attractant concentrations were calculated by multiplying the normalized average fluorescence intensity by the source attractant concentration. For high-magnification images (FRET), the average red-channel fluorescence was obtained from a 101 × 3 pixel region surrounding the cell centroid at each time point.

#### FRET measurements

Ratiometric measurements of RaichuCdc42 were corrected for non-FRET components by evaluating the coefficients of spectral bleed-through and cross-excitation, as described previously(Kamino *et al*., 2017). Namely, RaichuCdc42 expressing HL-60 cells were excited with 445 nm laser, and the fluorescence intensity at 528.5 –555.5 nm (IFRET) and 457–487 nm (ICFP) were obtained sequentially. The background-subtracted mean fluorescence intensities of the cell masks: ICFP and IFRET were used to obtain the “FRET index” ICFP/IFRET-corr as the final output. Here, the expected true FRET signal (IFRET-corr) was reverse-calculated from IFRET = IFRET-corr +α ξ ICFP + β ξ IYFP-dir. IYFP-dir is the fluorescent intensities at 528.5 –555.5 nm obtained by excitation at 488 nm laser taken independently per experimental run before timelapse acquisition. To obtain the coefficient for the spectral bleed-through (α) in our optical setup, cells expressing untagged CFP (mTurquoise GL) were illuminated with a 445 nm laser to obtain fluorescence intensities at the above two emission spectra (457–487 nm) and (528.5–555.5 nm). The average background fluorescence intensities obtained from 30 cell-free regions were subtracted before the coefficients were calculated. The coefficient of cross-excitation (β) was obtained by illuminating cells expressing untagged YFP (YPET) with either a 445- or 488-nm laser and recording the average fluorescence intensities at 528.5–555.5 nm. After background correction, the ratio between the two were taken to obtain β. To evaluate the spatial profile of Cdc42 activity, we calculated the weighted average of FRET signal ‘<FRET>’ along the reference coordinate vectors taken from the centroid origin; forward [y+] – backward [y-], cell front – rear, and the control axis [x+]-[x-] per cell masks (see equations in Supplemental Figure S2, B-D). To calculate these quantities, we first separated the cell mask into two sides: one side included the pixels located in the direction of the reference vector relative to the centroid, and the other included the pixels in the opposite direction. The fold change of Cdc42 activity was quantified from dividing <FRET> by its pre-stimulus average from t = -1 to 0 (min).

## Acknowledgements

The authors thank the present and past members of Sawai Lab for their insights and suggestions during this project. This work was supported by JSPS KAKENHI Grant Numbers JP15K12138, JP17H01812, JP17H05992, JP19H05801, and JP19KK0282 to SS; JP18J13059 to MI;

JP22H05673 to MU; and JST CREST JPMJCR1923 to SS and partly by JSPS KAKENHI Grant Number JP16K18537 to AN and HFSP Research Grant RGP0051/2021 to SS.

