## Supplemental Figures for "Traveling wave chemotaxis of neutrophil-like HL-60 cells"

Fig. S1

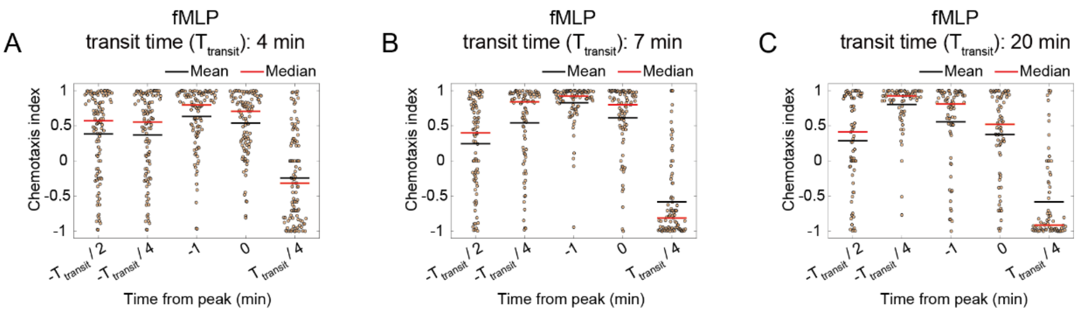


**Supplemental Figure S1:** Chemotactic index of cells at each stage of the fMLP wave The scatter dot plot (orange) indicates the chemotactic index of individual cells, with the mean (black) and median (red). Each panel shows the data for wave transit times of 4 min (A), 7 min (B), or 20 min (C).

Fig. S2

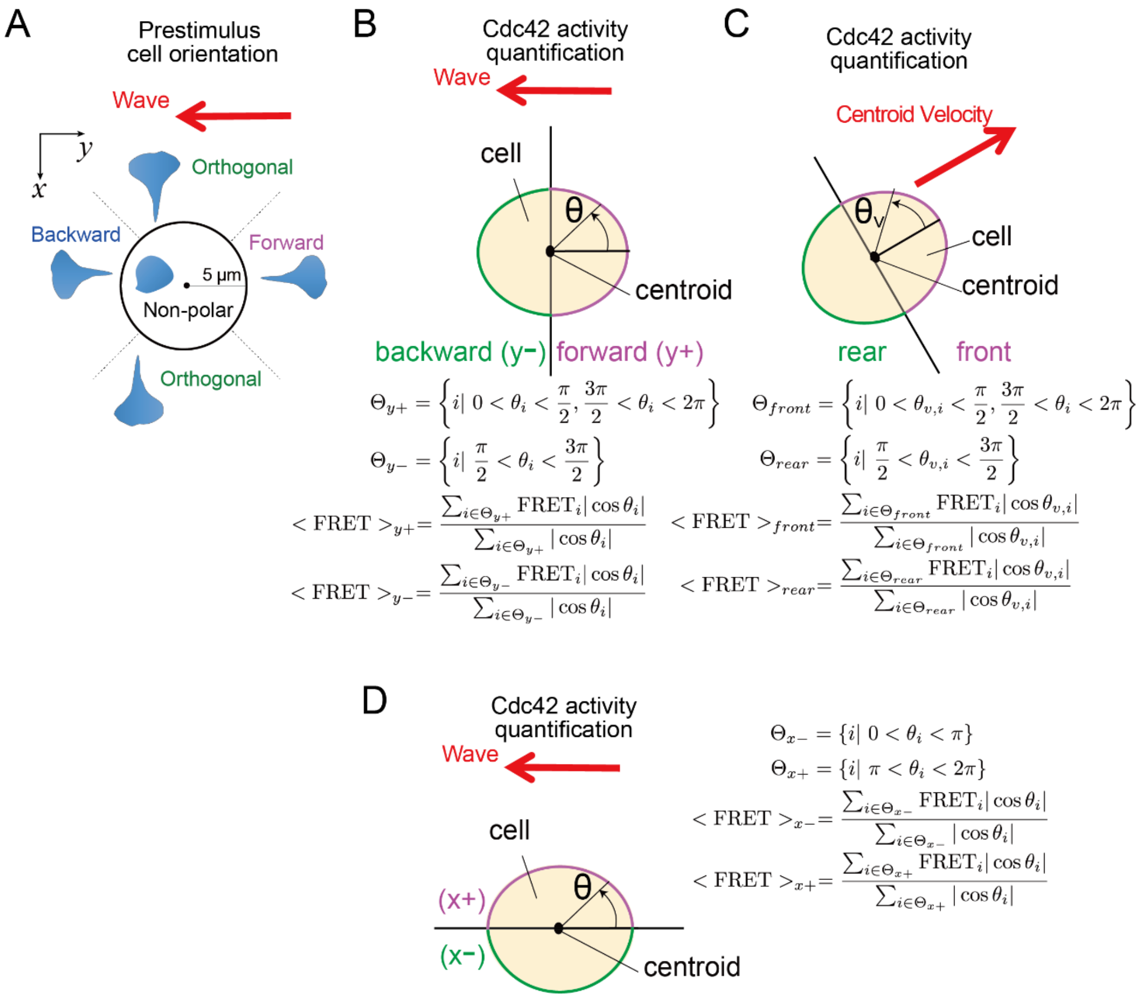


**Supplemental Figure S2:** Schematic of the analysis. (A) Pre-stimulus orientation (Forward, Backward, Orthogonal) or its lack (non-polar) with reference to wave propagation. (B-D) Cdc42-Raichu FRET quantification. The cell masks are divided into two hemispheres along the y-axis (B) based on the cell polarity axis defined by the centroid velocity vector (C) or along the x-axis (D). <FRET> of each hemisphere is calculated as the average of FRET where each pixel is weighted with the distance from the boundary between two hemispheres.

Fig. S3

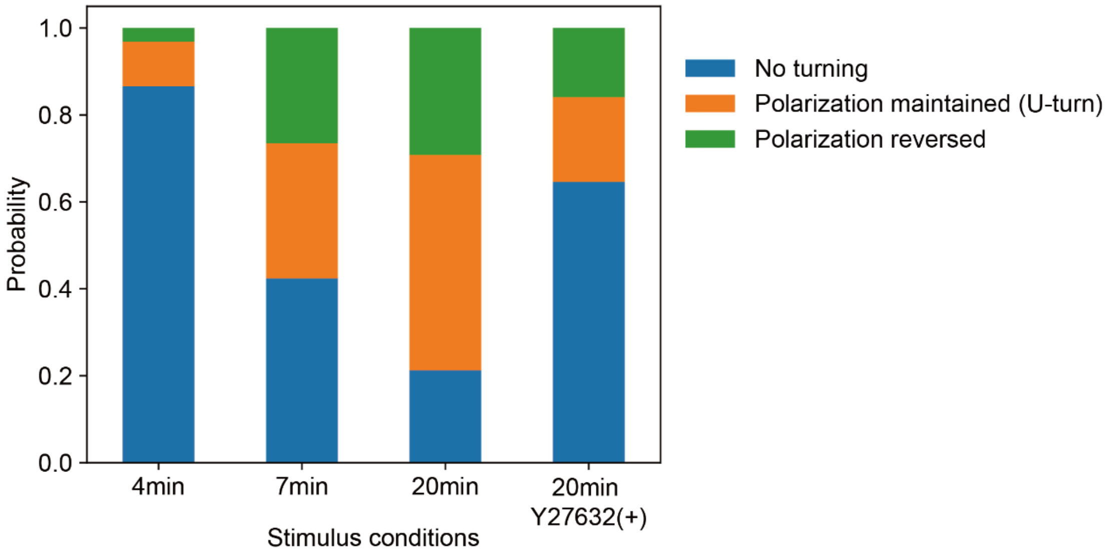


**Supplemental Figure S3:** Occurrence of turning behavior during waveback. Individual cell tracks are counted according to whether the cell exhibits no turn (blue: "No turning"), re-oriented while maintaining the cell polarity (orange: "Polarization maintained (U-turn)"), or re-oriented by producing a new leading edge (green: "Polarization reversed"). From left to right: Wave transit times of 4, 7, and 20 min for untreated cells, and 20 min for Y-27632-treated cells. Data are from those in Figures 1 and 9.

Fig. S4

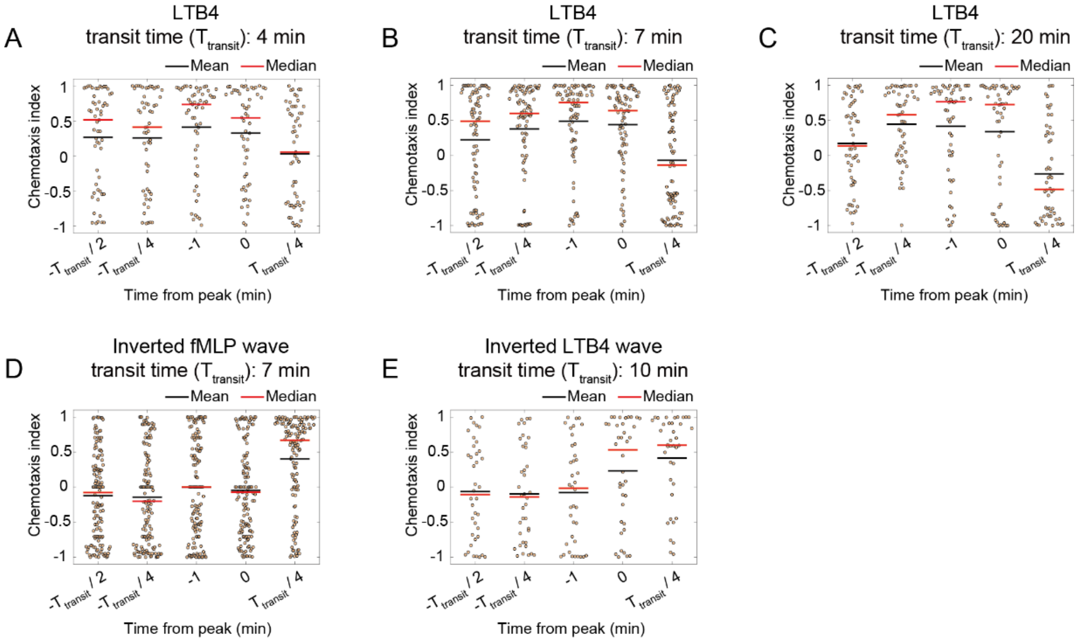


**Supplemental Figure S4:** Chemotactic index at each stage of the LTB4 wave or inverted wave of fMLP and LTB4. Scatter dot plot (orange) indicates the chemotactic index of individual cells, with the mean (black) and median (red). Chemotactic indices of cells for the (A-C) LTB4 wave, (D) inverted fMLP wave, or (E) inverted LTB4 wave. (A-C) Each panel shows the data for wave transit times of 4 min (A), 7 min (B), or 20 min (C). For the inverted waves (D and E), the transit times are 7 min (D) and 10 min (E).

Supplemental Fig. S5

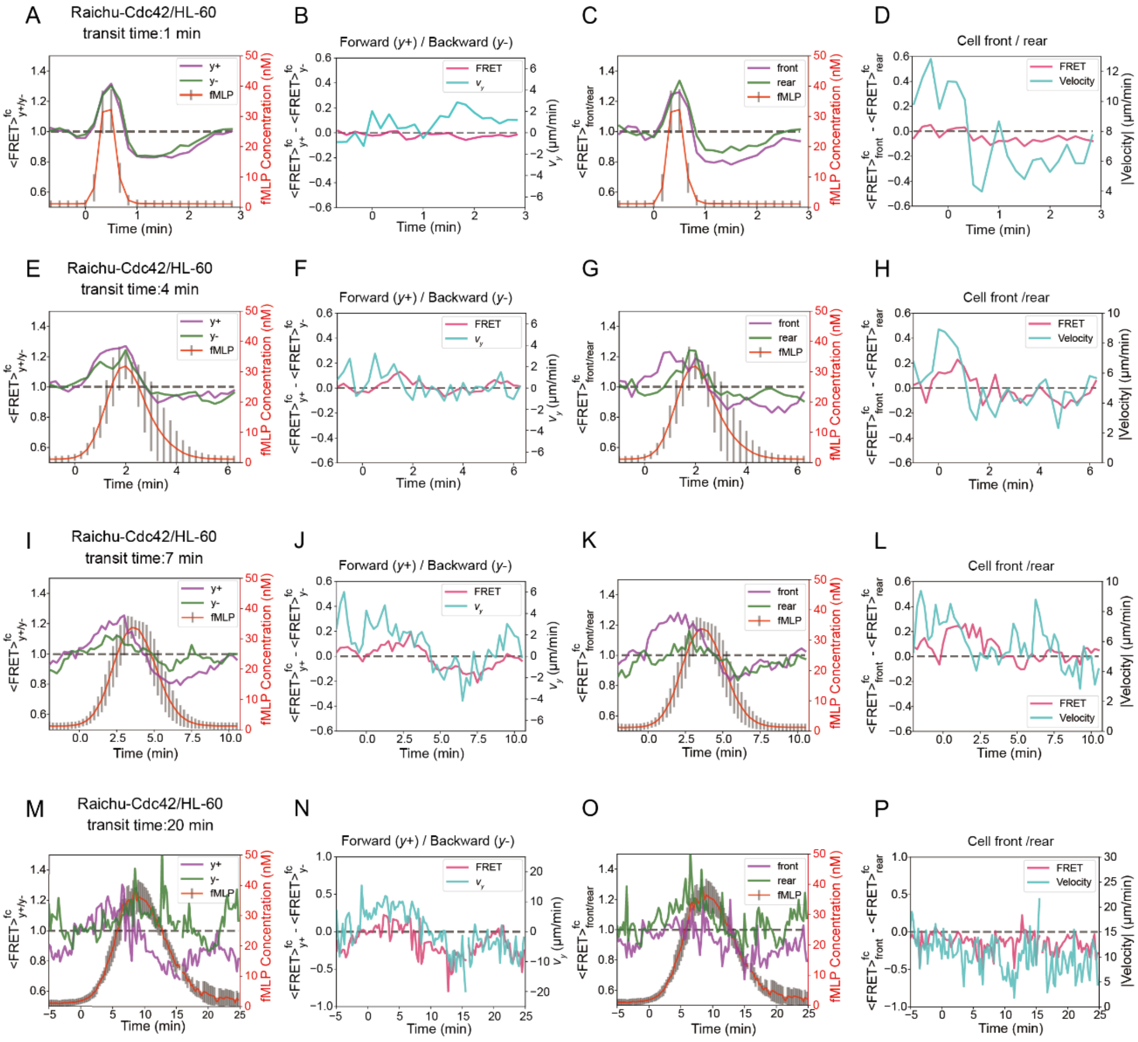


**Supplemental Figure S5: Cdc42 activity is a sum of basal activity and stimulus-dependent fold-change response.**

Fold-change in mean <FRET> for wave transit time of 1 min (A-D), 4 min (E-H), 7 min (I-L), and 20 min (M-P). The mean of normalized $\langle FRET \rangle_{y+}$ (magenta) and $\langle FRET \rangle_{y-}$ (green) (A, E, I, and M). The mean of normalized $\langle FRET \rangle_{y+}$ subtracted by normalized $\langle FRET \rangle_{y-}$ (magenta) plotted against y-velocity (cyan). (B, F, J, and N). The mean of normalized $\langle FRET \rangle_{front}$ (magenta) and $\langle FRET \rangle_{rear}$ (green) along the cell velocity vector (C, G, K, and O). The mean of normalized $\langle FRET \rangle_{front}$ subtracted by normalized $\langle FRET \rangle_{rear}$ (magenta) plotted against the averaged absolute speed (cyan) (D, H, L, and P).

Fig. S6

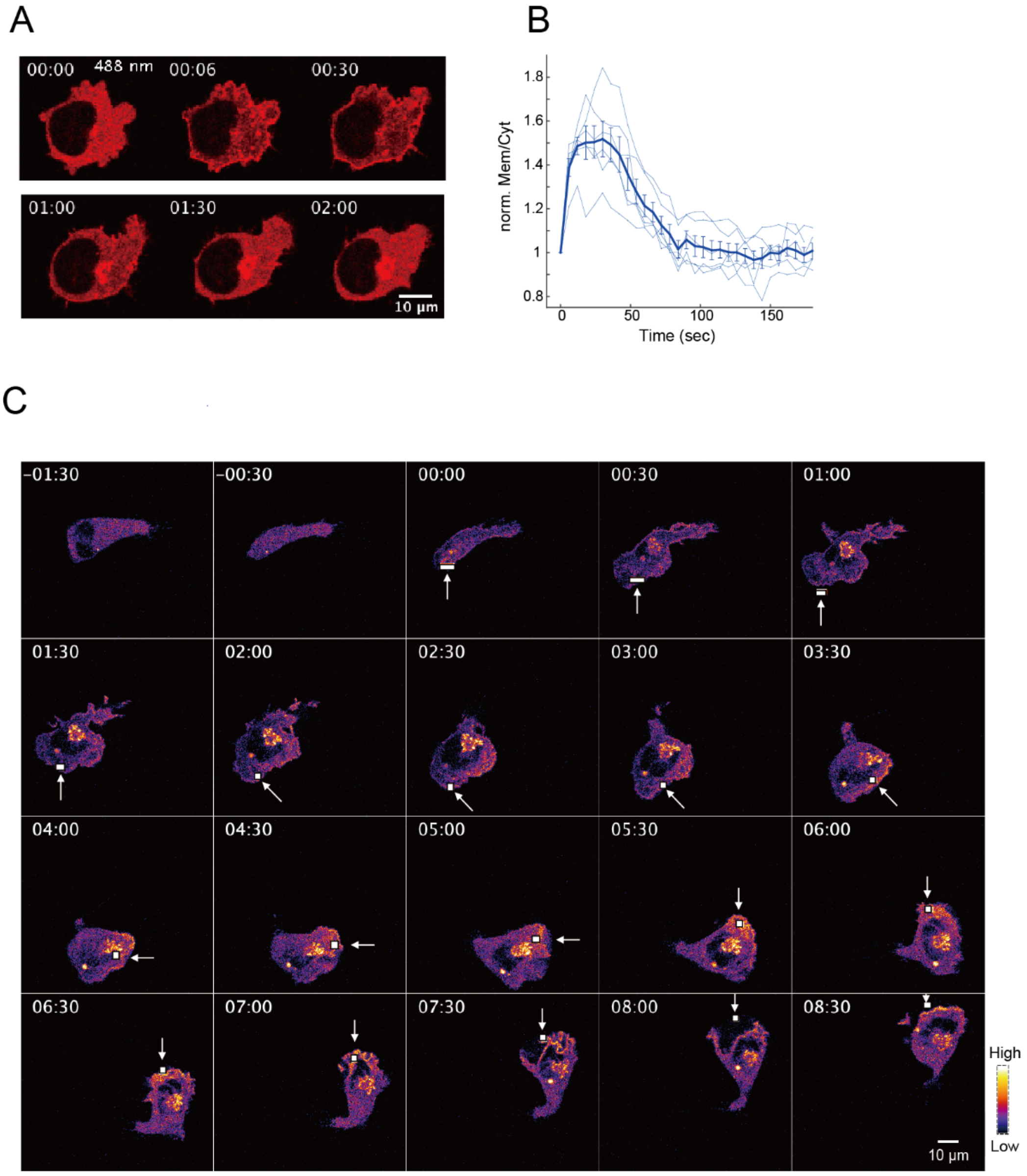


**Supplemental Figure S6:** Light-induced translocation of ITSN(DH/PH) induces a leading edge and orients HL-60 cells. (A and B) Time course of the average magnitude of light-induced translocation. Error bars indicate the standard error of the mean (n = 6 cells). Uniform stimulation induces transient localization that lasts for approximately 60 s. Snapshots (A) and mean fluorescence changes along the cell edge (B). The localization of ITSN (DH/PH) is quantified as the ratio of the mean fluorescence intensity in the membrane region to that in the cytosolic region of the cell mask. The ratio is normalized to the value in the time frame of blue light illumination. (C) Time series of successive applications in a cyclic trajectory and the resulting rotational cell movement. The arrows and white boxes indicate the light-illuminated regions. Representative data from 15 trials.

Fig. S7

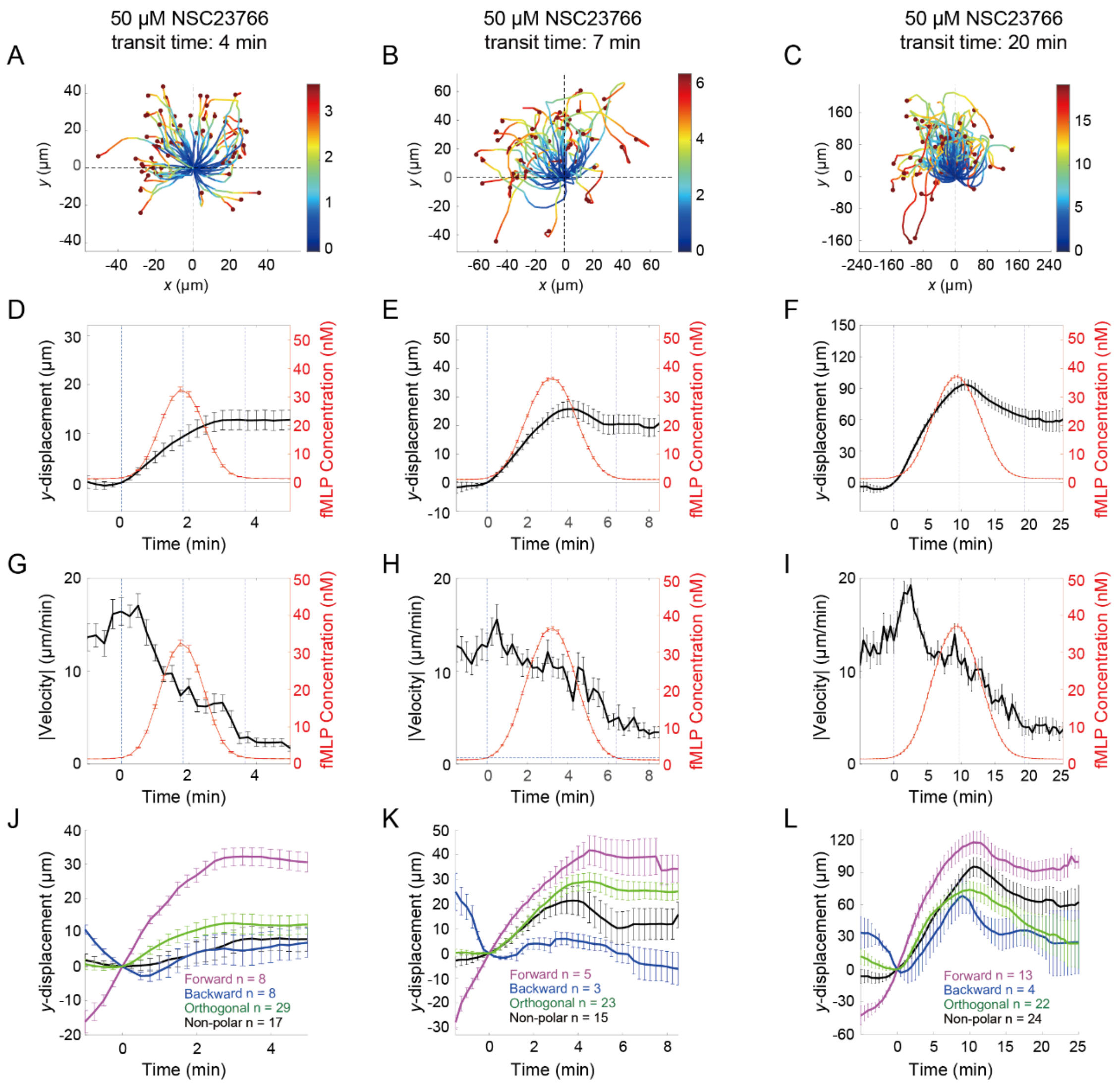


**Supplemental Figure S7:** Traveling wave chemotaxis in cells treated with the Rac inhibitor NSC23766. (A-C) Cell centroid trajectories for wave transit times of 4 (A), 7 (B), and 20 min (C). The origin is set to the initial position at the time of the stimulus arrival (t = 0). (D-F) Average centroid displacement in the y-direction (black) in reference to the LTB4 level (red) for wave transit times of 4 min (D), 7 min (E), and 20 min (F). Average magnitude of the centroid velocity for wave transit times of 4 min (G), 7 min (H), and 20 min (I). n = 62 (D and G), 46 (E and H) and 72 cells (F and I). (J-L) Mean y-displacement of cell centroids based on the initial cell orientation for wave transit times of 4 (J), 7 (K), and 20 min (L). Forward (magenta), backward (blue), orthogonal (green), and non-polar (black); see Supplemental Figure S2A for notations.

Fig. S8

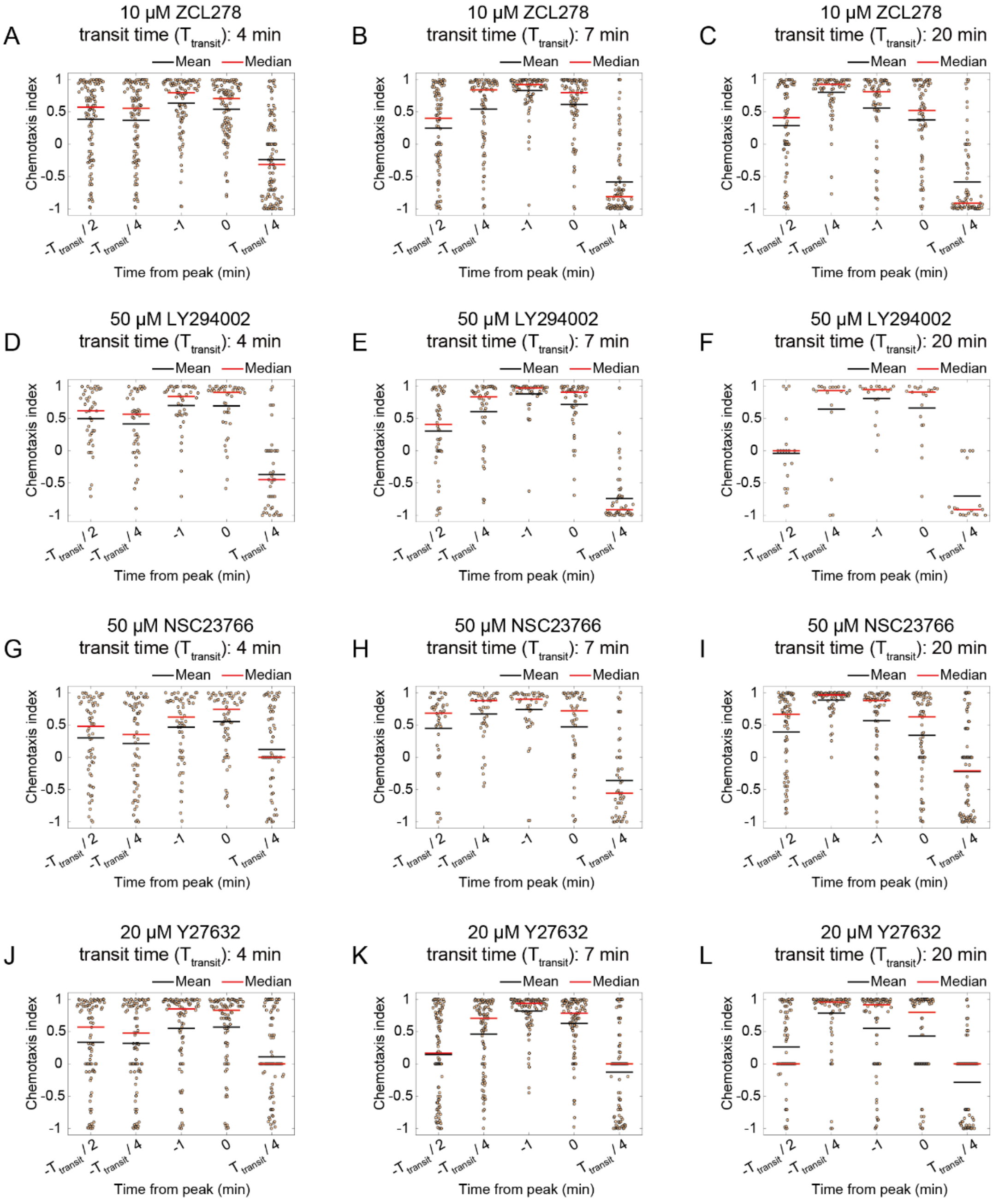


**Supplemental Figure S8:** Chemotactic index of inhibitor-treated cells at each stage of the fMLP wave. Scatter dot plot (orange) indicates the chemotactic index of individual cells with the mean (black) and median (red). Chemotactic index of cells for wave (A-C) with 10 µM ZCL278, (D-F) 50 µM LY294002, (G-I) 50 µM NSC23766, or (J-L) 20 µM Y-27632. Each column shows data for wave transit times of 4 min (A, D, G, and J), 7 min (B, E, H, and K), and 20 min (C, F, I, and L).
